# Integrative dissection of gene regulatory elements at base resolution

**DOI:** 10.1101/2022.10.05.511030

**Authors:** Zeyu Chen, Nauman Javed, Molly Moore, Jingyi Wu, Michael Vinyard, Luca Pinello, Fadi J. Najm, Bradley E. Bernstein

**Author notes:** These authors contributed equally (Zeyu Chen, Nauman Javed).

## Abstract

Although vast numbers of putative gene regulatory elements have been cataloged, the sequence motifs and individual bases that underlie their functions remain largely unknown. Here we combine epigenetic perturbations, base editing, and deep learning models to dissect regulatory sequences within the exemplar immune locus encoding CD69. Focusing on a differentially accessible and acetylated upstream enhancer, we find that the complementary strategies converge on a ∼170 base interval as critical for CD69 induction in stimulated Jurkat T cells. We pinpoint individual cytosine to thymine base edits that markedly reduce element accessibility and acetylation, with corresponding reduction of CD69 expression. The most potent base edits may be explained by their effect on binding competition between the transcriptional activator GATA3 and the repressor BHLHE40. Systematic analysis of GATA and bHLH/Ebox motifs suggests that interplay between these factors plays a general role in rapid T cell transcriptional responses. Our study provides a framework for parsing gene regulatory elements in their endogenous chromatin contexts and identifying operative artificial variants.

**Highlights:**

- Base editing screens and deep learning pinpoint sequences and single bases affecting immune gene expression
- An artificial C-to-T variant in a regulatory element suppresses CD69 expression by altering the balance of transcription factor binding
- Competition between GATA3 and BHLHE40 regulates inducible immune genes and T cell states

## Introduction

Genome-wide maps of chromatin state and transcription factor (TF) binding have nominated more than a million cell type-specific regulatory elements (REs) in the human genome as potential context-specific regulators of gene expression(Andersson and Sandelin, 2020; ENCODE Project Consortium et al., 2020; Stunnenberg et al., 2016). A critical next step is to determine their functions and sequence determinants. Computational tools that predict functional bases and/or gene targets are rapidly evolving, but require systematic benchmarking against perturbational data (Avsec et al., 2021; Nasser et al., 2021). Massively parallel reporter assays (MPRA) enable high-throughput analysis of sequence determinants within REs, but are based on exogenously introduced constructs that do not recapitulate the native chromatin contexts (Kheradpour et al., 2013; Klein et al., 2020; Maricque et al., 2018; Melnikov et al., 2012). CRISPR interference (CRISPRi) with fusions between dCas9 and the KRAB repressor provides a means to suppress a regulatory element in its native context and evaluate consequent transcriptional changes (Canver et al., 2015; Fulco et al., 2016; Gilbert et al., 2013; Korkmaz et al., 2016; Sanjana et al., 2016). Traditional CRISPR-based genetic perturbations offer increased resolution(Diao et al., 2017; Rajagopal et al., 2016), but may incur variable sequence changes due to heterogeneity of indels after DNA repair. Base editors can incur single base variants without frame-shifts or indels. They have been used to systematically characterize coding variants(Cuella-Martin et al., 2021; Gaudelli et al., 2017; Hanna et al., 2021; Kim et al., 2017; Komor et al., 2016), but have yet to be applied to noncoding REs.

In this study, we integrated CRISPRi, dCas9 and base editing with computational predictions to parse non-coding regulatory sequences in the CD69 locus. We found a ∼170bp interval within a ∼1500bp enhancer proximal to the CD69 promoter which plays a key role in regulating gene expression. Within this interval, base editing and deep learning converge upon a critical cytosine at chr12:9764948 (hg38), where a C-to-T transition significantly reduces element accessibility and CD69 expression. We show that this C-to-T base edit ablates a GATA3 binding site, thereby exposing a nearby E-box/bHLH site for BHLHE40 binding. Systematic analysis of chromatin accessibility and TF binding during T-cell activation supports a global role for binding competition between GATA3 and BHLHE40 in immune gene responses and T cell polarization.

## Results

### Resolving functional bases within immune regulatory elements

To dissect functional sequences within regulatory elements, we established a workflow combining chromatin profiling, deep learning, CRISPRi, dCas9 and base editing (**Figure 1A**). We combined ATAC-Seq accessibility maps with deep learning models to predict REs and functional sequences that regulate inducible gene expression in T cells. We then incorporated CRISPRi, dCas9 interference and base editing to directly test the regulatory functions of sequences and individual bases (Figure 1A).

**Figure 1.**
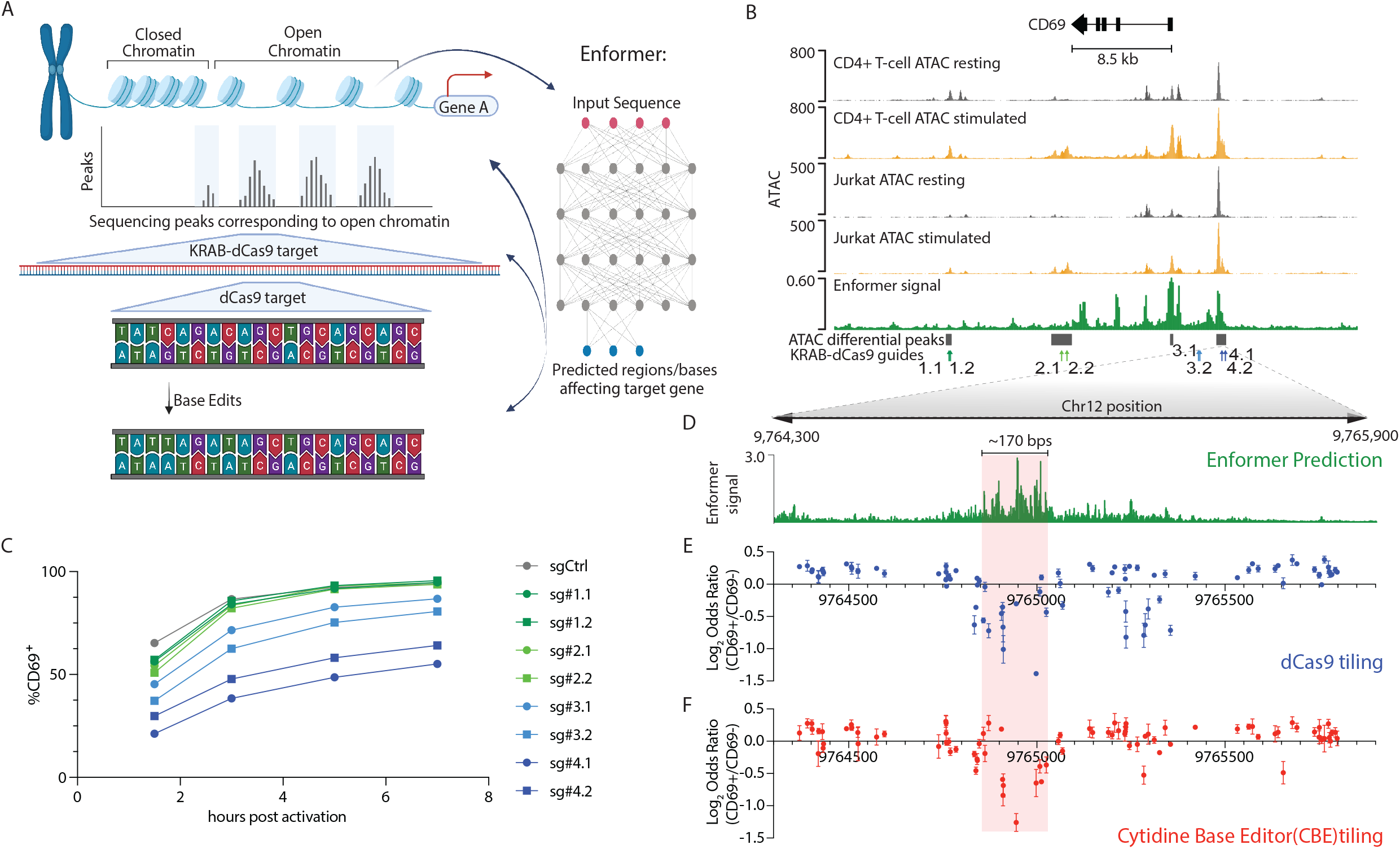
Integrative analysis of the CD69 regulatory landscape. A) Gene regulatory landscape characterization by successive functional assays and deep learning. B) Genomic tracks depict accessibility of the CD69 locus in primary CD4+ T cells and Jurkat cells, without or with stimulation (PMA/ionomycin). Enformer signal track shows the predicted contribution of underlying sequence to CD69 expression (magnitude of the model gradient at each position with respect to CD69 promoter signal, summed over 128 bp bins)in Jurkat. Grey bars depict regions with differential accessibility in stimulated Jurkat cells, relative to resting (FDR=0.2). CRISPRi sgRNA positions are also indicated. ATAC signal corresponds to reads per genomic content (RPGC). C) Flow cytometry of CD69 expression in Jurkat cells targeted with the indicated CRISPRi sgRNA following a stimulation time course. Samples gated from the lentiviral transduced population (mCherry+). D) Expanded view of Enformer signal at single base resolution over RE-4, as denoted in panel b. E) Enrichment/depletion plot of dCas9 sgRNAs in CD69+ Jurkat cells, relative to CD69-cells (y-axis; Log_2_ Odds Ratio of normalized sgRNA reads). sgRNAs along the x-axis according to their 5’ starting position on the positive strand. Each data point represents mean±s.e.m. F) Enrichment/depletion plot of Cytidine Base Editor (CBE) sgRNAs in CD69+ Jurkat cells, relative to CD69-cells (as in panel e). For C,E,F, data represent 2-3 biological independent experiments. A 170 bp region critical for CD69 activation is denoted (D-F, light red).

We focused on the CD69 locus, which encodes a key molecule for T cell signal transduction and tissue residency(Cibrián and Sánchez-Madrid, 2017; Sathaliyawala et al., 2013). CD69 expression is rapidly induced upon stimulation by T cell receptor cross linking or PMA/ionomycin in both CD4+ T cells and the Jurkat T cell line (**Figure S1A-S1B**). Chromatin accessibility maps nominated putative regulatory sites that gain accessibility upon stimulation in primary T cells and Jurkat cells (**Figure 1B**). We refined these predictions using the Enformer model (Avsec et al., 2021) trained on chromatin maps and CAGE-seq data (FANTOM Consortium and the RIKEN PMI and CLST (DGT) et al., 2014)(ENCODE Project Consortium et al., 2020). Genomic intervals corresponding to the promoter, 3’ UTR and an RE located ∼4 kb upstream of the TSS were predicted to impact CD69 transcriptional induction (**Figure 1B**).

We used CRISPRi to test the functional impact of the promoter (RE-3), the putative upstream RE (RE-4) and two other sites in the locus that also gained accessibility upon T cell activation (**Figure 1B and Table S1**). We infected Jurkat cells with lentiviral constructs containing KRAB-dCas9 and sgRNAs, selected positive cells, applied PMA/ionomycin stimulation, and measured CD69 surface protein expression by flow cytometry. We found that sgRNAs targeting RE-4 had the strongest suppressive effect on CD69 induction (**Figure 1C and S1C**), while sgRNAs targeting the TSS-proximal RE-3 had a weaker suppressive effect (**Figure 1C and S1C**). RE-4 corresponds to a DNase hypersensitive site bound by multiple TFs that has scored in a luciferase reporter assay and a CRISPR activation screen (ENCODE Project Consortium et al., 2020; Laguna et al., 2015; Mumbach et al., 2017). Whereas chromatin accessibility over RE-4 spans ∼1.4 kb, the Enformer model predicted that a specific ∼170 bp sequence interval within RE-4 is most critical for CD69 regulation (**Figure 1D and S1D**).

To resolve the functional sequences within these elements and test the Enformer prediction, we designed a library of 101 sgRNAs that tile sequences spanning RE-3 and RE-4 (**Figure S2A-S2B and Table S2**). We reasoned that dCas9 without the repressive KRAB domain would specifically occlude TFs overlapping its target site, and thus affect a narrower interval than KRAB-dCas9 (Dominguez et al., 2015). We infected Jurkat cells with a pooled lentiviral CRISPR library composed of dCas9 and the 101 sgRNAs, selected for puromycin resistance and stimulated with PMA/ionomycin for 5 hours. We then isolated genomic DNA from pre-sorted and sorted CD69- and CD69+ subsets (**Figure S2C**), and amplified the sgRNA-cassettes for sequencing. The relative effect of each sgRNA on CD69 expression was calculated based on its enrichment/depletion in CD69+ relative to CD69-libraries. Multiple sgRNAs within the ∼1.7 kb tiled region suppressed CD69 activation (**Figure 1E and S2D**).

To pinpoint individual functional bases in these REs, we complemented the dCas9 tiling with Cytidine Base Editor (CBE) and Adenine Base Editor(ABE) screens. We infected Jurkat cells with lentiviral constructs containing CBE or ABE and the same pool of 101 sgRNAs (**Figure. S2A-S2B, and Table S2**). We stimulated and sorted the cells, and then sequenced the sgRNA-cassettes from pre-sorted, CD69- and CD69+ subsets (**Figure S2C**). Multiple sgRNAs scored in these screens as reducing CD69 activation (**Figure 1F and S2E-S2G**). Notably, the CBE and dCas9 perturbations both pinpointed a ∼150 bp interval within RE4 centered at sg#70 as critical for CD69 expression (**Figure 2A**; Chr12:9764860-9765010). This experimentally identified interval closely coincided with the region identified by the deep learning model (**Figure 1D**). Several ABE hits in or near this interval also suppressed CD69 induction, but with lower fold-enrichment, potentially due to reduced effect sizes (**Figure S2G**).

**Figure 2.**
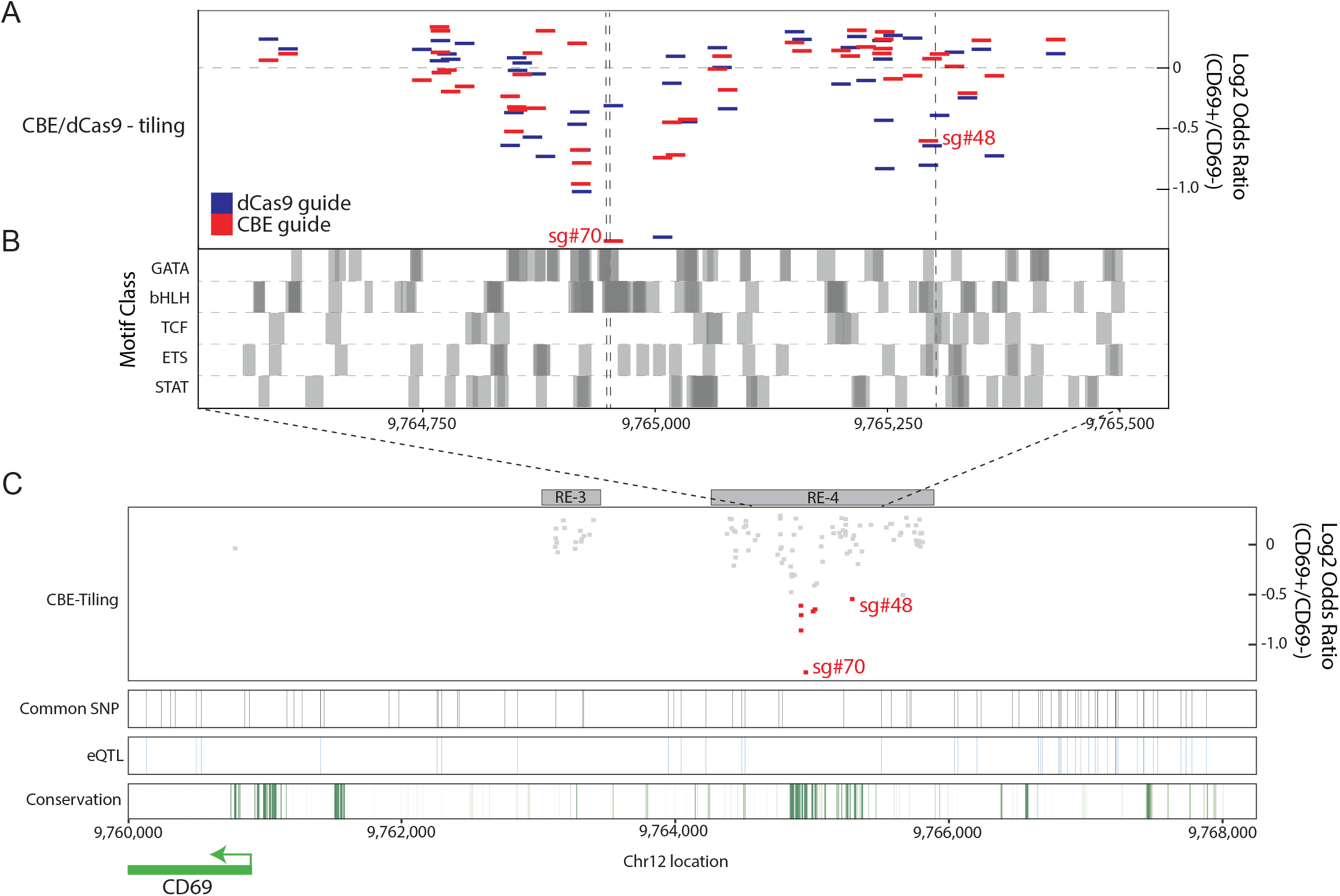
A critical sequence interval within RE-4 influences CD69 expression. A) Enrichment/depletion plot of sgRNAs in dCas9 and CBE tiling screens as in Fig 1e/f, limited to the central portion of RE-4 with sgRNAs shown to scale. Expected C->T edit positions highlighted for CBE-sgRNAs sg#70 and sg#48 (dashed grey lines). B) Transcription factor motif locations (grouped by broad motif class) for key immune regulators shown across the same interval as in panel a (FDR<0.05). Dark grey areas represent overlapping motifs. C) Zoomed out view of the CD69 locus shows CBE sgRNA depletion (red boxes indicate significantly depleted sgRNAs), common SNPs (black vertical stripes), eQTLs (blue vertical stripes)(Võsa et al., 2021) and PhastCon100 conservation score (green stripes).

The implicated interval in RE-4 is over-represented for multiple TF motifs relevant to immune function, including GATA, bHLH/Ebox, TCF, ETS and STAT (**Figure 2B**). Notably, a second top scoring interval from the CBE and dCas9 screens, centered at sg#48, showed similar TF motif enrichments (**Figure 2A**; Chr12:9765200-9765310). We scanned the locus for annotated expression quantitative trait loci (eQTLs). However, the implicated RE-4 intervals are highly conserved evolutionarily, devoid of natural variation in the human population, and thus invisible to eQTL analysis (**Figure 2C**)(Võsa et al., 2021). These findings highlight the importance of engineered variants for parsing highly conserved regulatory sequences.

### A single nucleotide artificial variant suppresses CD69 expression by affecting TF competition

We next sought to validate individual base edits and their transcriptional consequences. The top scoring CBE screen sgRNA, sg#70, is predicted to incur C->T transitions at positions 948 and/or 952 within RE-4 (chr12: 9,764,948 and 9,764,952). We infected Jurkat cells with a CBE vector containing either sg#70 or a control sgRNA (sgCtrl) and measured CD69 by flow cytometry (gating strategy in **Figure S3A**). This confirmed that CBE-sg#70 strongly reduced CD69 induction upon stimulation (**Figure 3A and S3B**). We next amplified and sequenced the target region from genomic DNA isolated from Jurkat cells infected with CBE-sg#70(Clement et al., 2019). In unsorted cells, C-948 was replaced by T on ∼57.0% of alleles. The proportion of C-948 edited alleles was higher in sorted CD69-Jurkat cells (67.0%) and lower in the CD69+ population (53.6%), consistent with a suppressive effect on CD69 induction (**Figure 3B**). In contrast, edits to the other candidate site, C-952, were less frequent (14.4% at baseline, 16.6% in CD69-, 13.9% in CD69+, **Figure 3B**). These results indicate that the single C-948->T edit strongly impacts transcriptional induction of CD69 in response to stimulation.

**Figure 3.**
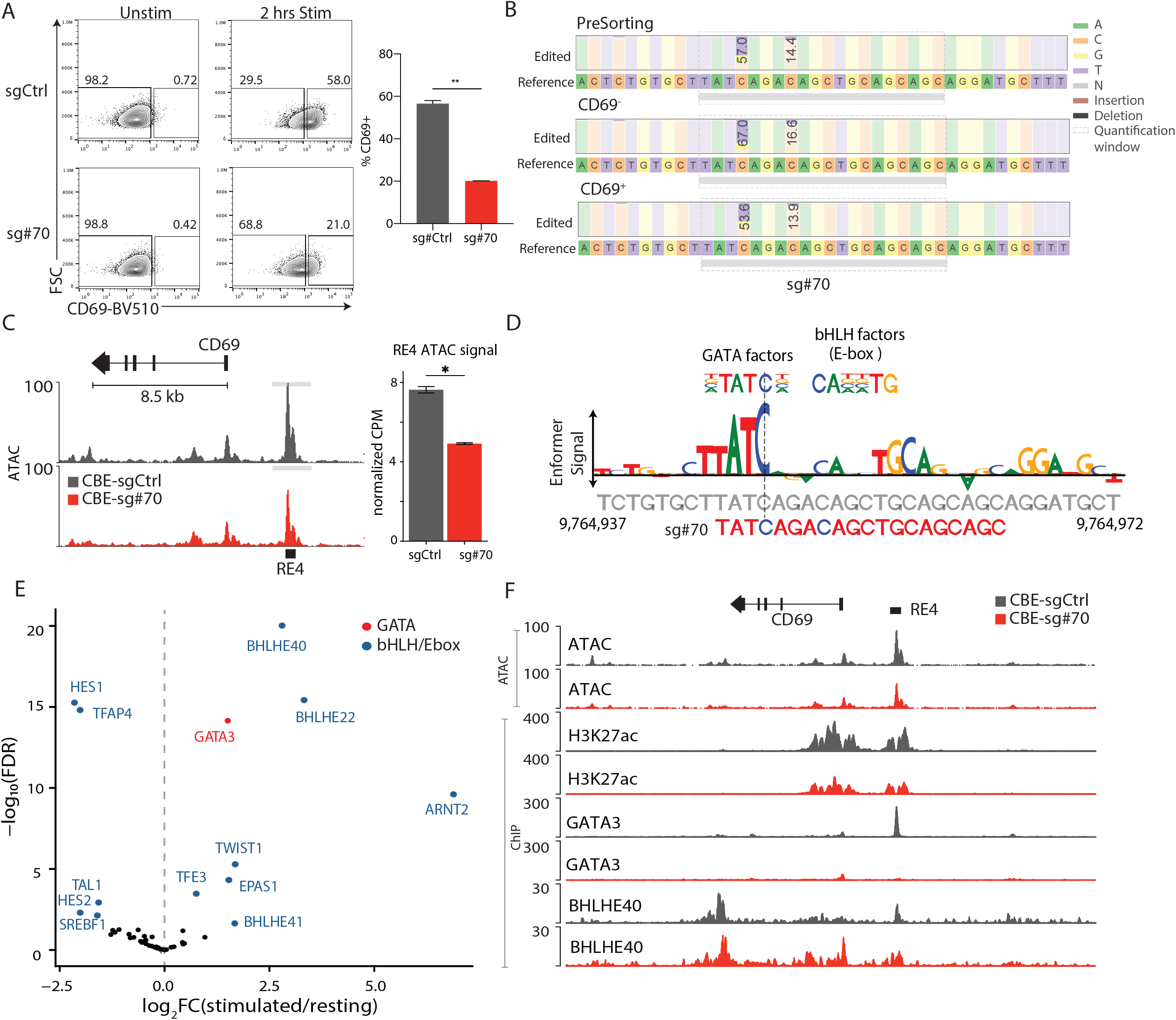
Top scoring base edits target competitive TF binding sites. A) Flow cytometry plots of CD69 signal for CBE-sgCtrl and CBE-sg#70 Jurkat cells under resting or stimulated conditions. Bar plot depicts the proportion of CD69+ cells in CBE-sgCtrl (grey) and CBE-sg#70 (red) after stimulation. P-value based on unpaired t test, **P<0.01. Data are from 4 independent experiments each with 2-3 technical replicates, mean±s.e.m. B) Table depicts frequency of incurred base edits in CBE-sg#70 infected Jurkat cells. PCR amplicons from unsorted, CD69- and CD69+ populations were sequenced by Illumina Nextseq500. Consensus sequence is shown along with stacked bars that depict the proportions of cytosine and thymine bases in the sequencing data (numbers indicate percent of alleles with C->T edit). Shaded boxes indicate the sg#70 target sequence. C) Chromatin accessibility shown over the CD69 locus for stimulated CBE-sg#70 (red) and CBE-sgCtrl (grey) Jurkat cells. Bar plot depicts the mean ATAC-seq signal over RE-4 (TMM normalized counts per million; CPM). P-value based on unpaired t test, *P<0.05. Data are from 3 replicates, mean±s.e.m. D) Enformer signal (letter height) for the sg#70 target region indicates the predicted impact of each base on RE-4 accessibility. The sgRNA directly coincides with a GATA motif and a bHLH/E-box motif, and incurs an edit that disrupts the former (vertical dashed line). E) Volcano plot depicts gene expression fold-change (x-axis) and significance (y-axis) for TF genes in stimulated Jurkat cells, relative to resting cells. Labels identify differential GATA (red) and bHLH/Ebox (blue) family members. F) Genomic tracks for the CD69 locus depict chromatin accessibility (ATAC), H3K27 acetylation (H3K27ac), GATA3 binding and BHLHE40 binding in CBE-sgCtrl (grey) and CBE-sg#70 (red) Jurkat cells. Y-axis represents the -log_10_(p-value) to input controls. Jurkat cells in a, b, c and f were stimulated with PMA/ionomycin for 2 hours.

We also examined the impact of the C-948 edit on chromatin accessibility. ATAC-seq profiles revealed reduced RE-4 accessibility in cells harboring the CBE-sg#70 construct, relative to CBE controls (**Figure 3C and S3C**). The reduced accessibility was specific to RE-4, as we did not observe any other accessibility changes in the CD69 locus or neighboring genomic regions (**Figure S3D**), nor in the vicinity of other activation associated genes such as CD28 and NR4A1 (**Figure S3E**). Hence, the single base substitution at position C-948 reduces RE-4 accessibility and suppresses CD69 induction in stimulated Jurkat cells.

We next considered the mechanism that underlies the potent effect of this single base mutation. Scanning the region for motifs showed that C-948 directly overlaps a GATA site predicted by the optimized Enformer model to impact both CD69 expression and element accessibility in Jurkat cells (**Methods; Figure 3D and S3F**). Importantly, the C-948->T edit disrupts a critical position in the GATA motif. The GATA motif is adjacent to a bHLH/Ebox motif that also scores in the Enformer model. We sought to identify specific TFs that are dynamically expressed and likely to bind these respective motifs (**Figure 3E**). GATA3 is highly expressed in Jurkat cells, up-regulated upon stimulation, and broadly implicated in T-cell lineage commitment (Ho et al., 2009). Among bHLH factors, BHLHE40 and BHLHE22 are both highly expressed and strongly induced upon stimulation. BHLHE40 in particular has established roles in T cell differentiation, inflammation and autoimmunity (Cook et al., 2020).

Whereas GATA3 is generally associated with transcriptional activation in T cells, BHLHE40 is a transcriptional repressor (Asanoma et al., 2015; Cook et al., 2020; Emming et al., 2020; Honma et al., 2002; Huynh et al., 2018; Zawel et al.). The close juxtaposition of their cognate motifs suggests that only one factor - either activator or repressor - can bind the implicated site at a given time. We therefore hypothesized that the paired motifs constitute a dynamic regulatory switch that contributes to CD69 induction. The potent suppressive effect of the base edit could then be explained by its ability to displace the GATA3 activator by disrupting its motif, allowing in turn BHLHE40 repressor binding due to relief of steric hindrance.

To investigate this hypothesis, we used ChIP-seq to map GATA3, BHLHE40 and the enhancer-associated histone acetylation mark H3K27Ac. Whereas a strong GATA3 binding peak is evident over RE-4 in stimulated Jurkat cells, binding is lost in CBE-sg#70 infected cells (**Figure 3F**). Remarkably, GATA3 loss in the edited cells is accompanied by broader BHLHE40 binding over RE-4, consistent with a switch in TF binding at the edited site (**Figure 3F**). H3K27ac signal over RE-4 is also reduced in the CBE-sg#70 edited cells, providing further support for the model that a switch from activator to repressor binding suppresses element activity (**Figure 3F**).

### BHLHE40 suppresses gene expression during Jurkat T cell activation via invade regulatory motifs near GATA binding sites

We further tested our model with GATA3 and BHLHE40 loss- and gain-of-function experiments. First, we confirmed that GATA3 knockout suppressed CD69 induction in stimulated Jurkat cells (**Figure S4A**). Next, we investigated GATA3-BHLHE40 antagonism by lentiviral BHLHE40 overexpression. We found that BHLHE40 overexpression suppressed CD69 induction in both control and CBE-sg#70 edited Jurkat cells (**Figure 4A**). However, the magnitude of suppression was greater in the edited cells, potentially due to relief of GATA factor competition. In contrast, overexpression of BHLHE41, the homolog of BHLHE40, had no effect on CD69 expression (**Figure S4B**). We also assessed the impact of BHLHE40 overexpression on chromatin accessibility using ATAC-seq. We found that overexpression reduced RE-4 accessibility, consistent with a direct repressive impact on the element and with our proposed TF switch model (**Figure 4B**).

**Figure 4.**
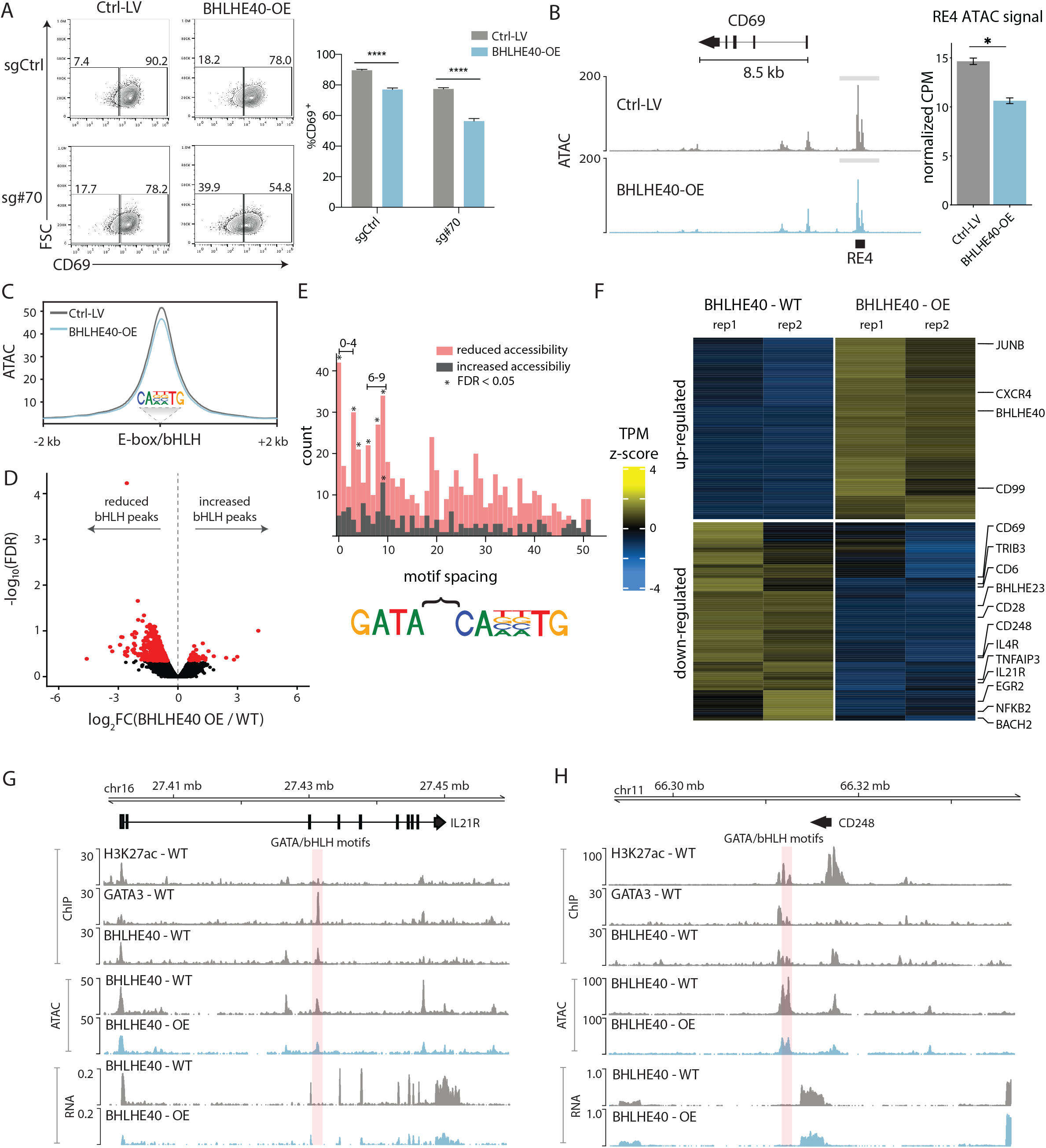
GATA3-BHLHE40 competition impacts global T cell transcriptional responses. A) Flow cytometry plots of CD69 signal for stimulated Jurkat cells transduced with CBE-sg#70 and a BHLHE40 overexpression construct (BHLHE40-OE), or with corresponding controls (sgCtrl and Ctrl-LV, respectively). Bar plot depicts the proportion of CD69+ cells in each condition. P-value based on unpaired t test, ****P<0.0001. Data are from 3 independent experiments with 2-3 technical replicates, mean±s.e.m. B) Chromatin accessibility in the CD69 locus for CBE-sg#70 Jurkat cells transduced with either BHLHE40 overexpression lentivirus (light blue) or control (grey). Cells were stimulated with PMA/ionomycin. P-value based on unpaired t test without multiple testing correction, *P<0.05. Bar plot data are from 2 replicates, mean±s.e.m ATAC-seq signal over RE-4 (TMM normalized CPM). C) Plot depicts aggregate accessibility (y-axis) for GATA3 bound sites that also harbor bHLH/E-box motifs (centered on the motifs). Data shown for stimulated Jurkat cells transduced with either BHLHE40 overexpression lentivirus (light blue) or control (grey). D) For the set of GATA3 bound sites with bHLH/E-box motifs in c, volcano plot depicts fold-change (x-axis) and significance (y-axis) of chromatin accessibility in Jurkat cells transduced with BHLHE40 overexpression lentivirus, relative to control. Differentially accessible sites (FDR < 0.1) are indicated in red. E) For differentially accessible sites in d, histogram shows the number of sites (y-axis) with the indicated spacing (x-axis) between GATA and bHLH/E-box motifs. Sites are stratified by whether their accessibility is reduced (red) or increased (grey) in the BHLHE40 overexpressing cells. Sites with significant peak differential between reduced and increased accessibility (FDR < 0.05) are denoted (*). F) Heatmap shows differentially expressed genes with BHLHE40 binding in their REs in BHLHE40 overexpressing Jurkat cells, relative to control. Cells were stimulated with PMA/ionomycin. G-H) Genomic views of the IL21R(F) and CD248(G) loci show ChIP-seq data for H3K27Ac, BHLHE40, and GATA3 in stimulated Jurkat cells (signal corresponds to P-value enrichment over input). Accessibility (ATAC) and expression (RNA-seq) are also shown for BHLHE40 overexpressing Jurkat cells and controls. Sites with combined GATA and bHLH/Ebox motifs are indicated (pink shade). Jurkat cells in b-f were stimulated with PMA/ionomycin for 2 hours.

Further evidence for the importance of interplay between these TFs emerged in our examination of the second interval identified in our dCas9 and CBE screens. Remarkably, the top base edit hit in this interval (sg#48) also incurs a C->T edit that disrupts a GATA motif flanked by a bHLH/Ebox motif (**Figure 2A-2B**). Here again, the respective motifs are too close to permit concurrent binding. Hence, this second hit may also be explained by its impact on competitive binding dynamics between the GATA3 and BHLHE40. This result prompted us to examine whether interplay between these factors plays a more general role in T cell transcriptional responses. We collated all GATA3 bound sites in Jurkat cells that contain a GATA motif and a bHLH/Ebox motif within the corresponding accessible site. We found that BHLHE40 overexpression reduced the aggregate accessibility of these sites (**Figure 4C-4D**), consistent with a global repressive role. Overall, 909 (95.4%) of these GATA3 sites with proximate bHLH/Ebox motifs became less accessible upon BHLHE40 overexpression, while just 44 (4.6%) became more accessible (FDR < 0.2). We also examined the edge-edge distance between the GATA and bHLH/Ebox motifs (**Figure 4E**). Sites that were repressed by BHLHE40 overexpression showed a strong enrichment for motif spacing of 0 to 3 bp, consistent with steric hindrance and competition between factors (FDR < 0.05). In contrast, a similar analysis of sites that were not repressed revealed a preferential spacing of 6 to 9 bp between motifs, consistent with previously reported sites of coordinate GATA and Ebox factor (e.g., TAL1) binding(Sanda et al., 2012). These findings suggest that the precise spacing of competitive or collaborative TF binding motifs is a critical determinant of regulatory element dynamics.

Finally, we considered the influence of the competitive TFs on T cell phenotypes. GATA3 is an established T cell regulator that promotes Th2 over Th1 differentiation(Wan, 2014), and is also implicated in the maintenance of naive T cells (Singer et al., 2017). Although BHLHE40 has also been associated with Th2 responses, we found that BHLHE40 overexpression downregulated multiple immune gene targets involved in Th2 differentiation (IL4R, IL21R, EGR2, etc.) or naive T cell maintenance (CD248, BACH2, etc.)(**Figure 4F**). Gene Set Enrichment Analysis confirmed that BHLHE40 overexpression upregulated Th1 response genes and effector T cell signatures, while down regulating genes associated with Th2 responses or naive T cells(Godec et al., 2016) (**Figure S5**). These results are consistent with a general role of BHLHE40 in restraining GATA3 mediated activation at immune loci. Notably, many of the immune loci subject to opposing regulation contain elements with closely spaced GATA and bHLH/Ebox motifs (0-3 bp), consistent with a general role for competitive TF binding on T cell transcriptional programs and phenotypes (**Figure 4G-4H**). We suggest that competition between repressor and activator poises key immune genes for rapid transcriptional responses, potentially explaining the association of both activator and repressor with T cell stimulation and Th2 phenotypes.

## Discussion

Resolving functional sequences within the vast numbers of putative regulatory elements in the human genome is a critical challenge with exciting potential to unlock an underlying regulatory code. Here we integrate chromatin maps, deep learning, epigenetic editing and base editing to parse sequences that control an exemplar inducible gene in Jurkat T cells. Regulatory base edits clustered in an evolutionarily conserved interval within a CD69 enhancer that was also highlighted by the deep learning model, but more precisely pinpointed critical regulatory sequences. Further characterization of top scoring edits revealed a role for competition between the GATA3 activator and the BHLHE40 repressor in the activation of CD69 upon T cell stimulation. Genomewide analysis suggests that dynamic interplay between these factors plays a general role in immune gene responsiveness and T cell phenotypes. Our study and results emphasize the importance of epigenetic perturbations and artificial sequence variants for characterizing regulatory sequences, which tend to be highly conserved and may be invisible to methods that rely on natural genetic variation.

We also note limitations of our study and approach. Although our pooled screen tested thousands of perturbations, it was limited to one inducible gene locus in one cell model. Extension of the approach to additional immune loci and in primary T cells is an exciting future opportunity. Furthermore, our base editing screen could target only ∼12% of nucleotide positions due to the requirement for nearby PAM sites. Critical bases and functional motifs will be missed as a consequence. Base editors with less restrictive PAM site requirements (Rosello et al., 2022) could improve the resolution of future screens. While base editor approaches are mainly focused on C-to-T or A-to-G transitions, prime editors could enable more systematic base changes if they could be applied at scale (Anzalone et al., 2019).

There remains a considerable gap between the throughput of current approaches and the eventual goal of deciphering the regulatory code of the entire human genome. Functional perturbations will need to be combined with computational approaches, such as the deep learning model incorporated here. While the model predictions and experimental data both highlighted similar genomic intervals in our study, the computational approach did not distinguish individual bases or motifs identified by the base editing. Nonetheless, algorithmic improvements, ideally trained in iterative cycles with experimental tests of artificial variants, may ultimately yield sufficiently accurate predictive models to resolve regulatory sequences across the vast noncoding genome.

Our study also highlights competitive interplay between GATA3 and BHLHE40 in the rapid induction of CD69 upon T cell stimulation. Two top scoring C-T base edits that suppress the CD69 response both appear to act by shifting the balance of TF binding from the GATA3 activator to the BHLHE40 repressor. Both cytosine bases and their surrounding regions are highly conserved, invariant in the human population, and hence invisible to QTL mapping studies. Hence, the artificial variants were essential to uncover functional bases, motifs and TF interactions. Interplay between GATA3 and BHLHE40 appears to play a much broader role in poising immune genes for T cell stimulation, with BHLHE40 repressing hundreds of GATA3-bound elements. The precise spacing between the GATA and BHLH/E-box motifs appears critical, with the antagonistic pairs tending to be very closely spaced, consistent with steric hindrance between TFs. Other adjacent motif pairs with wider spacing conducive to concurrent binding may have distinct biochemical properties and regulatory impacts. Thus our study links the well established principles of competitive and cooperative TF binding to specific motifs, functional elements, transcriptional responses and T cell phenotypes.

In conclusion, we have benchmarked emerging experimental and computational strategies to resolve regulatory genomic sequences with increasing precision. Our study demonstrates in particular the potential of base editing screens to identify critical regulatory motifs and TF interactions that underlie rapid and robust transcriptional responses. Further computational and experimental innovations will be needed to scale these approaches and address the daunting challenge of human regulatory genomics.

## Star Methods

### KEY RESOURCES TABLE

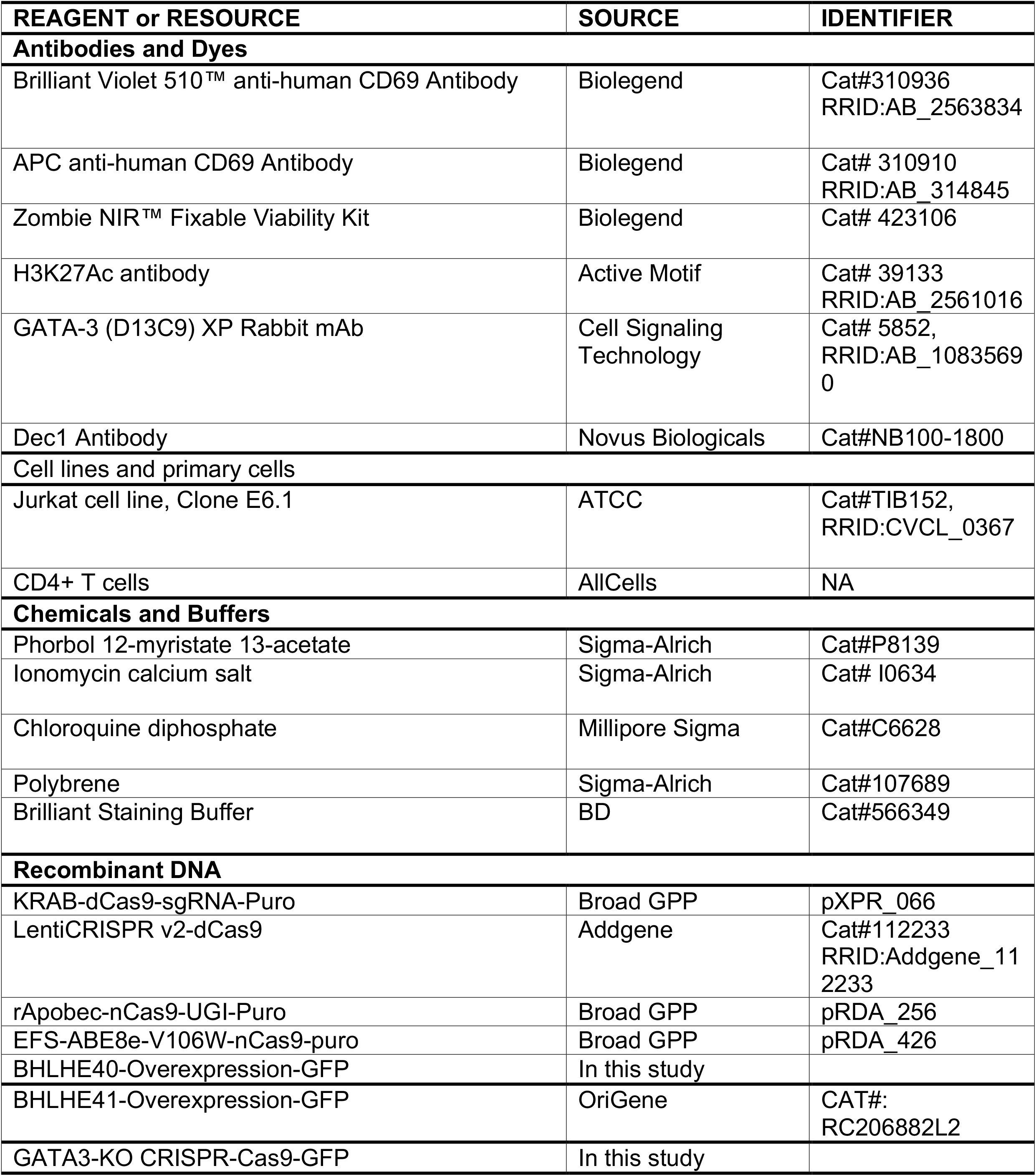
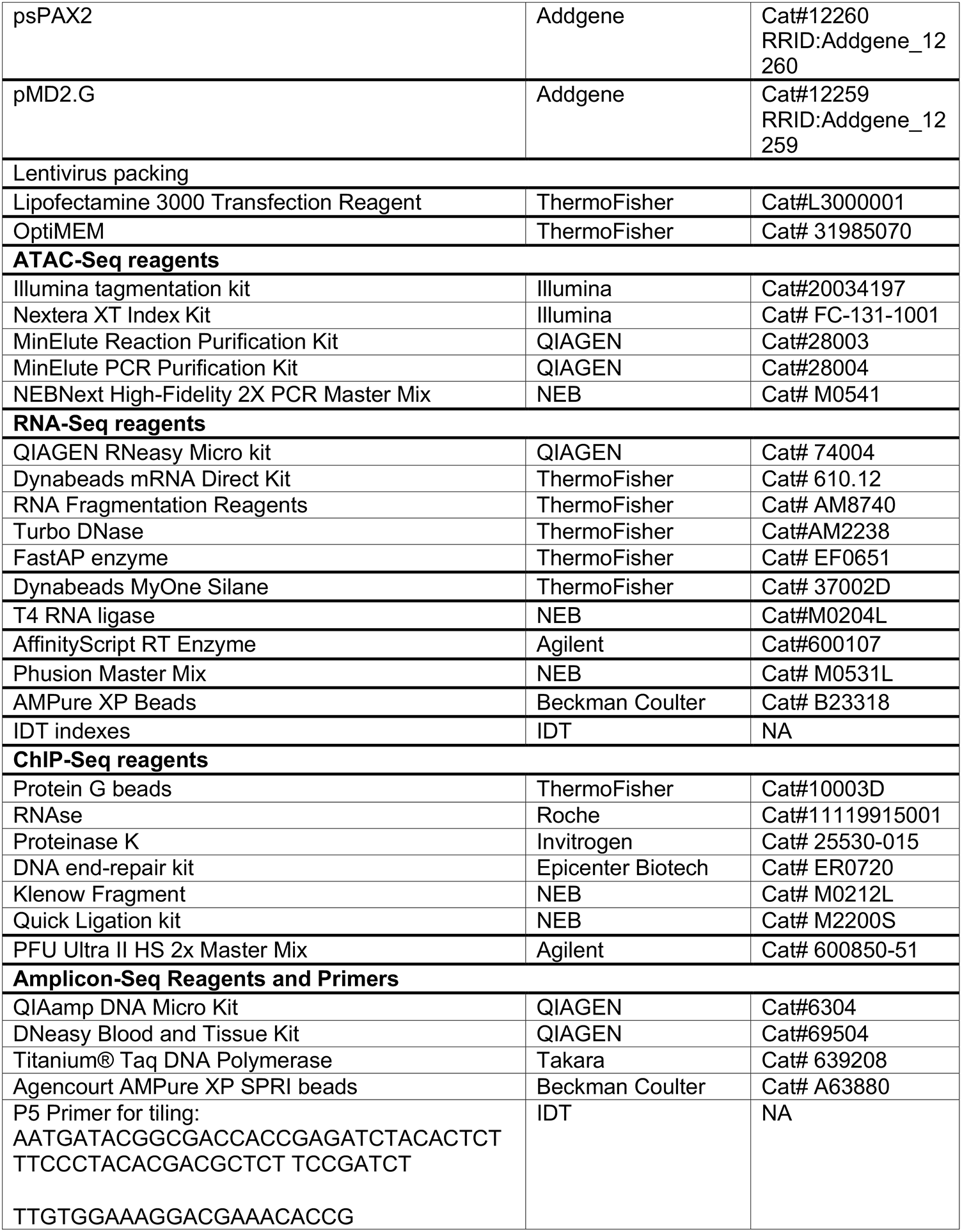
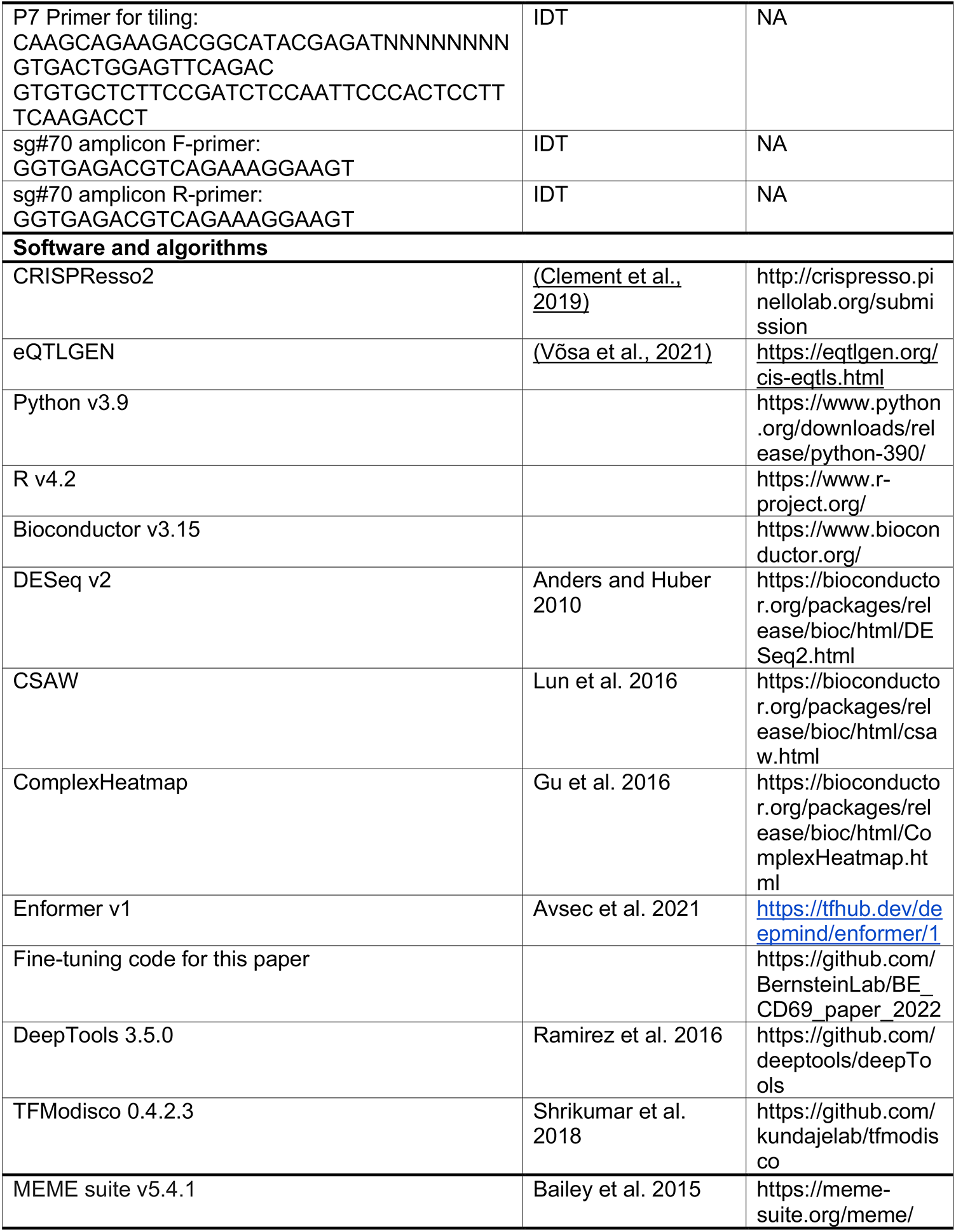
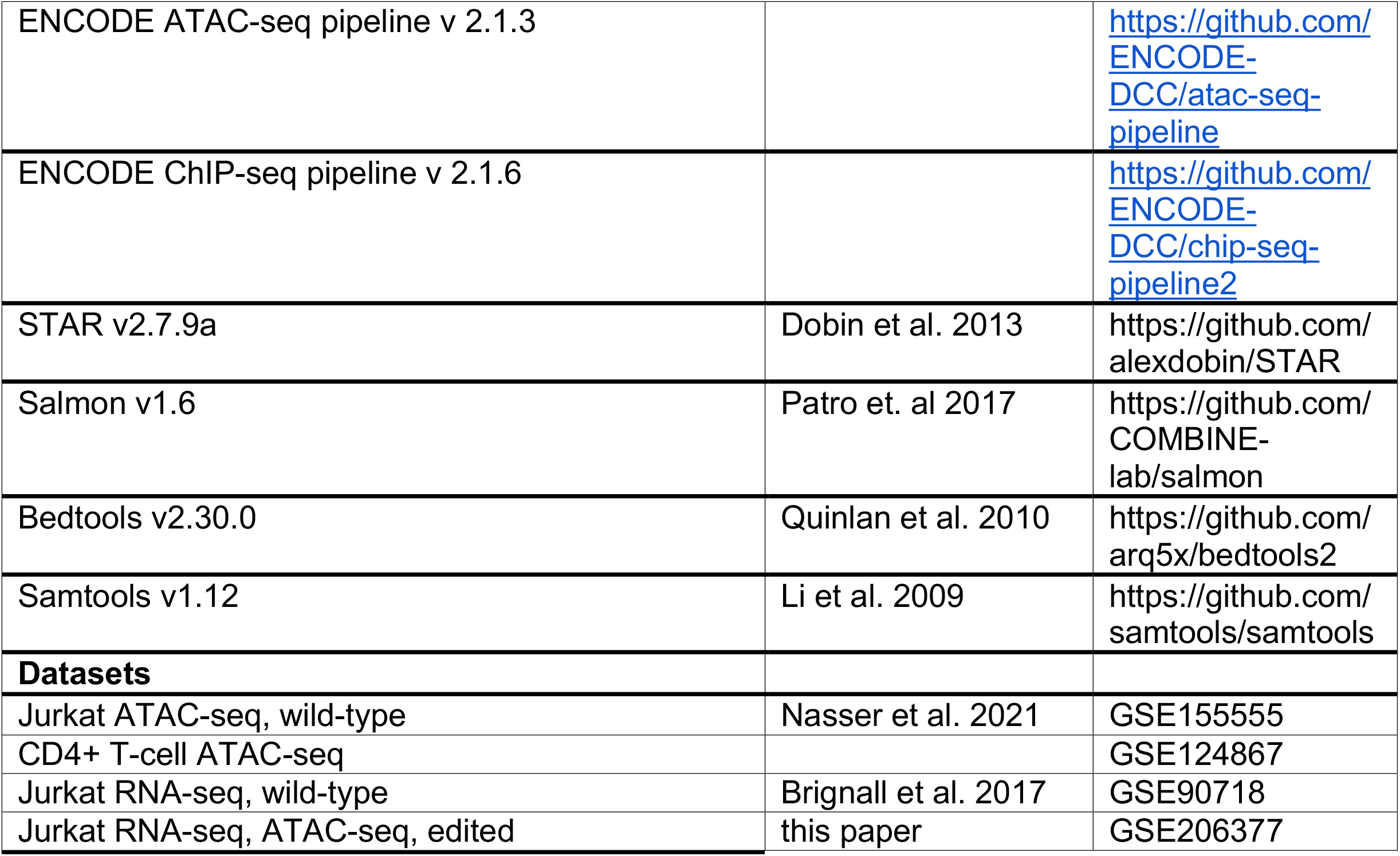

## Resource Availability

### Lead contact

**Further information and requests for resources and reagents should be directed to and will be fulfilled by the lead contact, Bradley E. Bernstein**.

## Material Availability

Base-editor construct will be available on addgene upon publication. sgRNA library request will be directed to Fadi J. Najm

## Data and code accessibility

For Jurkat and CD4+ T cell ATAC-seq datasets, we adapted GSE155555<u>(Nasser et al., 2021)</u> and GSE124867 for accessibility analysis on the CD69 loci. For the wild-type Jurkat RNA-Seq, we adapted GSE90718. ATAC-Seq data for 1) CBE+sgCtrl and CBE+sg#70, 2) Ctrl-GFP and BHLHE40-GFP, 3) CBE+sg#70+Ctrl-GFP and CBE+sg#70+BHLHE40-GFP, as well as RNA-Seq data for Ctrl-GFP and BHLHE40-GFP were generated for this study and are available at GSE206377.

Analysis code and custom scripts are available on github at https://github.com/BernsteinLab/BE_CD69_paper_2022.git.

## Method Details

### Guide library design and cloning

Pooled libraries for expression of sgRNAs were generated as detailed previously (Joung et al Nature Protocol 2017). Briefly, DNA oligos were annealed into double stranded fragments with compatible overhangs and ligated into BsmBI sites into vectors. Vector backbones were CRISPRi+guide puro (pXPR_066, Broad GPP), lentiCRISPR v2-dCas9 (gift of Thomas Gilmore, Addgene 112233), rApobec-nCas9-UGI-puro (pRDA_256, Broad GPP) and EFS-ABE8e-V106W-nCas9-puro(pRDA_426, Broad GPP). Libraries were then transformed by electroporation into electrocompetent coli (Invitrogen) and spread onto bioassay plates. Bacterial colonies were harvested and isolated using the Plasmid Plus Midi Kit (Qiagen). Four putative regulatory, ATACseq accessible regions were identified near the CD69 locus. Peak proximity and acceptable on-target efficacy scores(Doench et al., 2016) determined sgRNA selection for the CRISPRi tests. After RE3 and RE4 were identified, all sgRNAs possible in these peak regions were selected and included for screening with dCas9 and base editors and can be found in Table S2.

### Cell culture and stimulation

The Jurkat cell line (ATCC, Clone E6.1, TIB152) was cultured in complete RPMI (RPMI Medium 1640,Gibco, 11875085, 1% Penicillin-Streptomycin, Gibco, 15140122, 10% Heat Inactivate Fetal Bovine Serum, Peak Serum, 20mM HEPES,Gibco,15630080,1% Sodium Pyruvate, Gibco, 11360070, and 1% NEAA,Gibco, 11140050) at a maximum density of 2 × 10^6^ cells/ml in 25 cm or 75 cm cell culture dishes. Stimulation of Jurkat cells for 2-7 hour experiments was achieved with 50ng/ml Phorbol 12-myristate 13-acetate (PMA, Sigma-Alrich, P8139) and 500ng/ml ionomycin calcium salt from Streptomyces conglobatus (ionomycin, Sigma-Alrich, I0634).

Cryopreserved CD4^+^ T cells isolated from healthy donors were obtained from AllCells. On the day of stimulation, cells were thawed in RPMI 1640 medium supplemented with 2mM L-glutamine and 50% FBS, counted and resuspended in TexMACS medium (Miltenyi Biotec) supplemented with 20 IU/mL human Interleukin-2 (IL-2) and 1% penicillin-streptomycin. Cells were seeded at 1 million cells per well in a 48-well plate. Cells were either left untreated or stimulated with 10 μL T Cell TransAct™, human (Miltenyi Biotec) via CD3 and CD28 for 24hrs.

### Lentivirus production

293T cells approaching 70-80% confluency in 10 cm cell culture dishes were used for packaging. Cells were pre-treated with 25 uM chloroquine diphosphate (Millipore Sigma, C6628) in 3 ml of complete DMEM (Gibco DMEM with 1% Penicillin-Streptomycin and 10% Heat Inactivate Fetal Bovine Serum) and incubate in the 37°C and 5% CO2 incubator for more than 30 minutes. Lipofectamine 3000 Transfection Reagent (ThermoFisher, L3000001) was used to deliver plasmids into 293T cells. Briefly, 15 ug lentiviral vector plasmid, 15 ug of psPAX2 and 5 ug pMD.G plasmid were vortexed with 40 ul P3000 reagent in 1.5 ml OptiMEM (ThermoFisher, 31985070). Then 40 ul Lipofectamine was added to 1.5 ml OptiMEM and briefly vortexed. The two OptiMEM solutions were combined and mixed well by vortexing for 30s and incubated at room temperature for at least 20 minutes. Carefully, the OptiMEM mixture was added dropwise to 293T cells and incubated in a 37°C and 5% CO_2_ incubator for 6 hours. Media were aspirated and replaced with 5ml of fresh complete RPMI. Lentiviral supernatant was harvested between 24 hours and 48 hours after transfection.

### Lentivirus tranduction

Jurkat cells were resuspended in 1ml media and seeded at a density of 2-5 × 10^5^ cells per well of a 12-well plate. Lentiviral supernatant was supplemented with 8 ug/ml polybrene (Sigma-Alrich) added to the Jurkat cells. The plate was then centrifuged at 2000xg, 32°C for 60mins. Cells were then incubated at 37°C and 5% CO_2_ overnight and changed into complete RPMI on the next day. For GFP+ or mCherry+ marked lentivirus, cells were sorted or analyzed 5 days after transfection via flow cytometry. For blasticidin selection, 5 ug/ml of blasticidin was added to the transduced cells and selected for 14 days. For puromycin selection, 5 ug/ml of puromycin was added to the transduced cells and selected for 3 days.

### Flow cytometry and sorting

Suspended cells were centrifuged down at 300xg, room temperature for 5 minutes. The cells were stained with the antibody cocktail in the staining buffer of a 1:1 mix of PBS and Brilliant Staining Buffer (BD, 566349), at room temperature for 20 mins or at 4°C for 30-40 mins. Cells were washed once in PBS with 1% FBS and then resuspended in the same buffer. Flow cytometry or FACS was processed on either BD LSRFortessa X-20 or SONY SH800 following the manufacturing instructions. Antibodies and dyes used from Biolegend: Brilliant Violet 510™ anti-human CD69 Antibody (310936);APC anti-human CD69 Antibody (310910); Zombie NIR™ Fixable Viability Kit (423106).

At least 2 × 10^5^ CRISPR library infected Jurkat cells were collected as a pre-sorted baseline. 2-4 × 10^6^ CRISPR library infected Jurkat cells were resuspended in 2 ml of complete RPMI and stimulated with 50 ng/ml PMA and 500 ng/ml ionomycin for 5 hours, and then processed for FACS as described above. Sorted CD69- and CD69+ populations were collected for genomic DNA isolation.

### Genomic DNA isolation and sequencing

Genomic DNA (gDNA) was isolated using QIAamp DNA Micro Kit (QIAGEN, 6304) or DNeasy Blood and Tissue Kit (QIAGEN, 69504) according to the manufacturer’s protocol. The gDNA concentrations were quantified by Qubit. For PCR amplification, at least 330 ng of gDNA was used per reaction for greater than 500-fold library coverage. Each reaction contained 1.5 ul Titanium Taq (Takara), 10 μl of 10× Titanium Taq buffer, 8 μl deoxyribonucleotide triphosphate provided with the enzyme, 5 μl DMSO, 0.5 μl P5 stagger primer mix (stock at 100 μM concentration), 10 μl of a uniquely barcoded P7 primer (stock at 5 μM concentration), and water up to 100ul.

P5 Primer: AATGATACGGCGACCACCGAGATCTACACTCTTTCCCTACACGACGCTCT TCCGATCT

TTGTGGAAAGGACGAAACACCG

P7 Primer: CAAGCAGAAGACGGCATACGAGATNNNNNNNNGTGACTGGAGTTCAGAC

GTGTGCTCTTCCGATCTCCAATTCCCACTCCTTTCAAGACCT

PCR cycling conditions included: an initial 5 min at 95°C; followed by 30 s at 54°C, 30 s at 53°C, 20 s at 72°C, for 28 cycles; and a final 10-min extension at 72°C. PCR primers were synthesized at Integrated DNA Technologies. PCR products were purified with Agencourt AMPure XP SPRI beads according to the manufacturer’s instructions (Beckman Coulter, A63880). Samples were sequenced on a MiSeq (Illumina). Reads were counted by alignment to a reference file of all possible guide RNAs present in the library. The read was then assigned to a condition on the basis of the 8-nt index included in the P7 primer.

### Amplicon sequencing

To assess base editing frequency of the sg#70 locus, we designed primers flanking this region resulting in a 214bp product. Forward primer: GGTGAGACGTCAGAAAGGAAGT and reverse primer: AATTCACCCACTGAAAGGAAAA. Amplicons were next ligated with Illumina Truseq adaptors, cleaned and size selected with AMPure XP SPRI beads, and sequenced on a MiSeq paired end run. FASTQ files were processed with CRISPResso2 v2 with standard settings for base editor(Clement et al., 2019).

### ATAC-Seq experimental processing

ATAC-Seq lysis buffer contains 10mM Tris-HCl (pH=7.4), 10mM NaCl, 3mM MgCl2, 0.1% Tween-20, 0.1% NP40, 0.1% Digitonin, 1% BSA and topped up with ddH2O. ATAC-Seq washing buffer contains 10mM Tris-HCl (pH=7.4), 10mM NaCl, 3mM MgCl2, 1% BSA and topped up with ddH2O.

5 × 10^4 cells were centrifuged down with the resuspension buffer (PBS with 1%BSA) in a low-binding eppendorf tube at 4°C, 500xg for 5 mins. Each pellet is resuspended with 50 ul of lysis buffer and incubated on ice for 5 minutes. 50 ul of wash buffer was added to the lysis buffer containing nuclei and centrifuged down at 4°C, 500xg for 5 minutes. The supernatant is then removed and 50ul resuspension buffer is added to the tube without disturbing the pellet. Nucleus are then centrifuged down at 4°C, 500xg for 5minutes. Tagmentation of the genome DNA is processed using the Illumina tagmentation kit (20034197) for 30 mins in 37°C. Fragmented products are then isolated via MinElute Reaction Purification Kit (QIAGEN, 28003) according to the manufacturer’s instructions. Illumina Nextera XT SetA indexes and NEBNext High-Fidelity 2X PCR Master mix (NEB, M0541) are used to amplify the fragmented products of each sample, with 12 PCR cycles of 98°C-10s, 63°C-30s and 72°C-1min. PCR products are then isolated via MinElute PCR Purification Kit (QIAGEN, 28004) following manufacturer’s instructions.

### RNA-Seq experimental processing

Whole RNA was extracted from over 1 × 10^5 cells using the QIAGEN RNeasy Micro kit (QIAGEN, 74004) according to the manufacturer’s instructions. 1ug RNA was then used to prepare the RNA-Seq library. Poly-A+ RNA is enriched using Dynabeads mRNA Direct Kit (ThermoFisher, 610.12) according to the manufacturer’s instructions and eluted in 18ul Tris-HCl buffer(pH=7.4). Zinc fragmentation are processed using RNA Fragmentation Reagents(ThermoFisher, AM8740), followed by Turbo DNase (ThermoFisher, AM2238) and FastAP enzyme (EF0651) treatment. Then the fragmented RNA are cleaned-up using Dynabeads MyOne Silane (ThermoFisher, 37002D) and eluted in 7ul of nuclease-free water. Next, RNA-adaptors are ligased to eluted RNA using T4 RNA ligase (NEB, M0204L) at 23°C for 1 hour and adaptor-ligated RNA was cleaned-up using Dynabeads MyOne Silane and eluted in 13.5ul of nuclease-free water. First strand of cDNA is synthesized using AffinityScript RT Enzyme (Agilent, 600107) according to the manufacturer’s instructions at 54°C for 1 hour. First-strand cDNA was cleaned-up using Dynabeads MyOne Silane and eluted in 5.5ul of nuclease-free water, followed by cDNA adaptor ligation. After another round of clean-up, the adaptor-ligated cDNA was processed to library PCR amplification using Phusion Master Mix (NEB, M0531L) with IDT adaptor indexes. The final library was cleaned-up with AMPure XP Beads (Beckman Coulter, B23318) to a final size around 280bps.

### ChIP-seq experimental processing

Jurkat cells were pelleted (2.5×10^7 per sample) and fixed using 1% formaldehyde at 37°C for 10 mins then quenched by glycine. Samples were next washed with cold PBS+proteinase inhibitor (ThermoFisher, 78429), resuspended in lysis buffer (1% SDS, 0.25% DOC, 50mM Tris-HCl, pH=7.4), and incubated on ice for 10 mins. Samples were diluted up to 1ml in eppendorf using ChIP dilution buffer (0.01% SDS, 150mM NaCl, 0.25% Triton, 50mM Tris-HCl, pH=7.4) and sonicated using a Covaris E220, with the following settings: 24 mins with 5% duty factor, 140W max power and 200 cycles/burst. Each sample was then split into 4 eppendorf tubes: 1) 20ul, top up to 200ul for input; 2)180ul, top up to 1ml for H3K27Ac ChIP (2.5ul, Active Motif, 39133); 3) 400ul, top up to 1ml for GATA3 ChIP (10ul, CST-D13C9, 5852); 4) 400ul, top up to 1ml for BHLHE40 ChIP (10ul, Novus Biological, NB100-1800). The tubes were incubated overnight at 4°C on a rotator.

On the next day, Protein G beads (ThermoFisher, 10003D) were washed and added to the antibody-containing suspension and rotated at 4°C for 2 hours. The beads were then washed with ice-cold RIPA wash buffer: RIPA-500, LiCl, and 10mM Tris-HCl buffer (pH=8.5). The beads were eluted in a wash buffer (10mM Tris-HCl, pH=8.0, 0.1% SDS, 150mM NaCl, 5mM DTT) and incubated at 65°C on a shaker for 1 hour. Samples were then treated with RNAse (Roche, 11119915001) at 37°C for 30 mins and then with proteinase K (Invitrogen, 25530-015) at 63°C for 3 hours. AMPure XP Beads (Beckman Coulter, B23318) were used to purify the DNA fragments from the samples. Eluted fragments were then processed for DNA end-repair (Epicenter Biotech, ER0720), Klenow A base adding (Klenow from NEB, M0212L), adaptor ligation (Ligase from NEB, M2200S) and PCR amplification (PFU Ultra II HS 2x master mix from Agilent, 600850-51) according to manufacturer’s protocols. Index primers were ordered from Integrative DNA Technology. PCR was set up with the following conditions: 2 mins for 95°C; 30 sec at 95°C, 30 sec at 55°C,30 sec at 72°C for 16 cycles; 1 min at 72°C. PCR products were purified using AMPure XP Beads with a final size of around 300 bps.

### Enformer predictions and fine-tuning

The published Enformer model without any modifications was downloaded from https://tfhub.dev/deepmind/enformer/1. For model fine-tuning, we loaded the model checkpoint made available by the authors at https://github.com/deepmind/deepmind-research/blob/master/enformer/enformer-training.ipynb. The cell-type/organism specific heads in the original model were then replaced with two untrained dense layers, corresponding to ATAC-seq from resting and stimulated Jurkat T-cells not in the original training data. These data were first converted to RPGC normalized bigwigs using DeepTools and then converted the required model input format using the scripts publicly available at https://github.com/calico/basenji. The modified model was then trained on a Google Cloud TPU-VM v3-64 pod-slice using a multi-learning rate scheme. The original model trunk, consisting of all convolutional and transformer layers shared for all organisms/tracks was trained using the AdamW optimizer from the tensorflow addons library at a learning rate of 1.0e-05 and weight decay of 1.0e-05. The two added output heads were trained at a higher learning rate of 5.0e-03 and weight decay of 5.0e-02. The model was trained for 52 epochs, with checkpointing every 8 epochs, and training was stopped when the validation loss did not decrease by 1.0e-03 relative to the lowest recorded validation loss for 30 epochs. The best checkpointed model was chosen at epoch 24 which reached a validation pearson’s correlation of 0.8021 and 0.7859 for stimulated and resting Jurkat T respectively.

Model interpretation was conducted as described at https://github.com/deepmind/deepmind-research/blob/master/enformer/enformer-usage.ipynb. For CAGE-seq interpretation, we calculated the gradient of the model for unstimulated Jurkat T-cells with respect to the predicted CAGE-seq signal at the CD69 promoter. This was achieved by centering a 393216 bp genomic window within the CD69 promoter(chr12:9760820-9760903) and computing the gradient for human output head # 4831 with respect to output bins 446-450.The absolute value of the gradients were then summed in 128bp bins for coarse grain resolution (**Fig 1D**). A similar approach to nominate bases contributing to RE-4 accessibility was adopted to obtain the base resolution contribution scores for the fine-tuned model corresponding to Figure S2 and 3. For this analysis, the window was centered around RE-4(chr12:9764300-9765900) and the gradient was computed with respect to output bins 442-454(**Fig S1D, 3D)**.

For identifying TF motifs using Enformer base importance scores (**Fig S3**), we used the TFModisco suite (Shrikumar et al., 2018). This tool clusters short stretches of bases using base importance scores to discover motifs that can then be matched to known databases. First we centered 393216 bp genomic windows as above at the promoter of each of 2195 genes that were differentially expressed(FDR=0.01, see RNA-seq processing below) between resting and stimulated Jurkat cells. Then, we computed the gradient of the model at each base within the window for output head 4831 as above with respect to the CAGE-seq signal at the promoter, corresponding to bins 446-450. For each window, we also computed the model gradient on a dinucleotide shuffled version of the sequence which was averaged across all genes in order to obtain an empirical null distribution of gradients. In order to reduce computing time, we extracted model gradients, sequence, and null gradients for the 750 centered bp window centered at each ATAC-seq peak detected from unstimulated Jurkat cells. Predictions were run in parallel across all genes simultaneously using a custom WDL/Google Cloud script. Finally, hypothetical contribution scores at each position within the 750 bp input window were computed as the model gradient corresponding to each non-reference base. TFmodisco was then used with default settings in order to identify putative regulatory motifs. Candidate seqlets were then matched to HOCOMOCO v11 motifs(Kulakovskiy et al., 2018) using Tomtom from the MEME-suite V5.4.1(Bailey et al., 2015).

### ATAC-seq data processing, differential accessibility analysis, and coverage tracks

All ATAC-seq data were aligned and processed using the ENCODE uniform ATAC-seq processing pipeline v2.1.3 available at https://github.com/ENCODE-DCC/atac-seq-pipeline. The pipeline was configured to use default parameters, adapter auto-detection, the bowtie2 aligner (Langmead and Salzberg, 2012), and MACS2 (Zhang et al., 2008) for peak calling. GRCh38 V29 and associated mitochondrial genomes and blacklists were obtained from https://www.encodeproject.org/references/ENCSR938RZZ/. Differential accessibility analysis was conducted with the CSAW package v 1.28(Lun and Smyth, 2016). Briefly, a consensus set of peaks was obtained from the union of peaks across all input samples/replicates. Peaks lying within blacklist regions and with low signal(less than -3 log_10_CPM) were removed. Reads were counted in 300 bp windows genomewide and merged to a maximum width of 5 kb. Finally, counts were TMM normalized and significant differentially accessible peaks were identified based on a genome-wide (for global analysis) or local window (<= 1 Mb) FDR correction.

ATAC-seq tracks in all figures were computed by pooling replicates where applicable using samtools (Li et al., 2009) and creating signal tracks using DeepTools bamcoverage (Ramírez et al., 2016). Signal tracks were normalized using the reads per genomic bin normalization (RPGC) options in DeepTools with the pre-computed effective genome size for GRCh38 in order to create coverage bigwigs. Average ATAC-seq coverage profiles over bHLH/E-box sites were obtained by taking genome-wide motif-scans (see motif analysis below) and filtering to keep bHLH(Ebox) motif sites overlapping the union of ATAC-seq peaks between the BHLHE40-OE and BHLHE40-WT conditions. DeepTools plotProfile function with a 4kb window centered at each bHLH(Ebox) motif was then used to compute the mean coverage profile for each condition.

### RNA-seq data processing, differential expression analysis, and gene-set enrichment

RNA-seq datasets were processed using a custom pipeline utilizing fastp v0.23.2((Chen et al., 2018) for automatic adapter trimming with default settings for paired-end datasets, and the STAR aligner (Dobin et al., 2013) v2.7.9a with default GTEx (GTEx Consortium, 2020) settings obtained from https://github.com/broadinstitute/gtex-pipeline. Gene quantifications were obtained using Salmon v1.6 (Patro et al., 2017) and the GENCODE V38 annotation (Frankish et al., 2021) with the seqBias, gcBias, posBias, and validateMappings flags enabled.

Differential expression analysis between conditions was conducted using DESeq V2 (Anders and Huber, 2010) with default settings. Log fold change values were corrected with the lfcShrink option using the apeglm method. For BHLHE40-OE gene expression analysis, BHLHE40-OE was compared to wild-type only in the stimulated case. Unless otherwise stated, significance is based on an FDR=0.05 cutoff.

Heatmap was constructed using the Complex heatmap package v 2.12.0 (Gu et al., 2016). Genes were subsetted to only keep those differentially expressed between BHLHE40-OE and BHLHE40-WT at FDR=0.05, and those with a nearby(< 50 kb peak-promoter distance) BHLHE40 ChIP-seq peak(see ChIP-seq processing below).

Gene-set enrichment analysis was conducted using the FGsea package v 1.22.0 (Korotkevich et al., 2021) and the ImmuneSigDB gene sets (Godec et al., 2016). The product of -log_10_(p-value) and log_2_FoldChange was used as the ranking metric for input to gene set enrichment. Significant pathways were collapsed using the collapsePathways function from FGsea with default settings. Normalized enrichment scores for enriched, down-regulated gene sets (identified as those containing the suffix _DN) were multiplied by -1.

### ChIP-seq data processing

ChIP-seq read alignment, quality filtering, duplicate marking and removal, peak calling, signal generation, and quality-control was conducted using the ENCODE ChIP-seq pipeline v2.1.6 available at https://github.com/ENCODE-DCC/chip-seq-pipeline2. GRCh38 V29 and blacklists were obtained from https://www.encodeproject.org/references/ENCSR938RZZ/. In brief, reads were aligned to the GRCh38 genome using bowtie2(-X2000), filtered to remove poor quality reads(Samtools) and de-duplicated(Picard MarkDuplicates). Both histone and TF peaks were then called using MACS2(Zhang et al., 2008). All datasets were assessed for enrichment quality and replicate concordance using the included ENCODE ChIP-seq quality control pipeline. H3K27ac ChIP-seq datasets passed all ENCODE QC standards and were not further processed beyond obtaining signal tracks as described below. For GATA3 ChIP-seq in the unedited cells (sgCtrl-P258), we noted a small number of peaks and therefore increased the MACS2 q-value cutoff to 0.05 and max number of peaks to 500000 to improve sensitivity. Consensus peak sets were then obtained using IDR analysis between replicates with an FDR cutoff of 0.05. De-novo motif discovery using the XSTREME (Grant and Bailey, 2021) program from the MEME-suite(Bailey et al., 2015) yielded the expected GATA motif among the top discovered motifs. A final peak set for sgCtrl-P258 was then obtained by retaining peaks that contained the expected GATA motif, and that overlapped open chromatin regions as defined by ATAC-seq. For BHLHE40-P258(sgCtrl), replicates showed poor concordance and the replicate(replicate 1) with the greater number of peaks was selected for further analysis. A peak set was obtained replicate 1 was obtained by taking the intersection of peaks between pseudo-replicates for this dataset. Motif discovery for this peak set yielded the expected e-box motif(CANNTG) among the top recovered motifs. As with GATA3, only peaks overlapping with ATAC-seq peaks and which contained an E-box/bHLH motif were retained. Target genes for each factor were then nominated by computing the set of genes with an annotated TSS within 50kb of a called peak using bedtools closest.

### Motif Analysis

To avoid motif redundancy, we obtained non-redundant motif cluster definitions and the corresponding PWMs from https://resources.altius.org/~jvierstra/projects/motif-clustering-v2.0beta/ (Vierstra et al., 2020). Motif scan of the RE-4 region corresponding to chr12: 9764556 - 9765505 was conducted by extracting the regions genomic sequence using bedtools getfasta (Quinlan and Hall, 2010) and scanned using the MOODs motif scanner v1.9.4.1 (Korhonen et al., 2009) with a p-value cutoff of 0.0001 and background base probabilities of 2.977e-01 2.023e-01 2.023e-01 2.977e-01. We filtered to keep motifs matched with MOODs score > 4, and further clustered motifs based on whether the motif cluster name contained GATA, bHLH, TCF, ETS, NFKB, NFAT, CREB, or STAT.

For motif spacing analysis, genome-wide motif scans were obtained from https://www.vierstra.org/resources/motif_clustering. Each GATA3 ChIP-seq peak(see ChIP-seq analysis) was intersected with the set of consensus ATAC-seq peaks for both the BHLHE40-OE and WT stimulated Jurkat samples. For the remaining peaks, the highest MOODs score GATA motif located within the middle ? of the peak was chosen as the representative GATA motif. The closest bHLH/E-box motif was then located using the bedtools closest tool with the -t all and -d options enabled. The motif-motif distance was then computed as the edge-edge distance between the central GATA motif and the core CANNTG E-box motif(i.e. the sequence GATAGGCACCTG would yield a distance of 2 bp). Motif pairs were then separated based on whether the ATAC-seq peak showed reduced or increased accessibility with BHLHE40-OE(FDR=0.2). Motif spacing enrichment was then computed by obtaining the number of motif-spacings at each distance from 1 to 50 bp for each group separately.Enrichment significance was tested based on the SpaMo approach (Whitington et al., 2011). For a given motif-pair with a maximum motif-motif distance of r, we assume that the number of sequences exhibiting a motif-motif distance 0 < x < r is binomially distributed, with all spacings being equally likely. This means that the probability of a given pair having a specific motif-spacing, ignoring strand and motif-orientation, is 1 / r. We choose restrict r <= 50, reasoning that motif pairs exceeding this edge-edge distance are unlikely to represent meaningful TF interactions, and to reduce the burden of multiple testing. The probability that we will observe a specific number of motif-pairs *n* out of a total number of motif-pairs *m* exhibiting a specific motif-spacing *x* by chance alone is then computed as 1.0 - CDF_binomial(n,m,1.0 / r). This p-value is then corrected for multiple testing using the BH procedure.

### Common SNP, eQTL and Conservation Score Analysis

Common SNPs are adapted from Ensembl GRCh38(Cunningham et al., 2022), with a cut-off of more than 1% minor allele frequency. Expression quantitative trait loci(eQTL) are adapted from eQTLGEN(Võsa et al., 2021), only cis-eQTLs are considered here with FDR< 0.05 (data from https://eqtlgen.org/cis-eqtls.html, gene locus for CD69). Conservation score are adapted via phastCons100way score(0-1, clear to dark green)(Siepel et al., 2005), sg#70 regions are marked specifically in the plot.

## Acknowledgment

We thank S. M. Bevil, S. Battaglia, G. Rahme, K. Macias and J. Verga for technical assistance. We thank the Google TPU Research Cloud for providing TPU access and support. This work was supported by funds from the NCI/NIH Director’s Fund (DP1CA216873 to B.E.B.), the Gene Regulation Observatory and the Variant-to-Function Initiative at the Broad Institute. Z.C. is supported by NCI-CA-234842. L.P. is supported by the National Human Genome Research Institute (NHGRI) (R35HG010717, UM1HG012010). J.W. is supported by a postdoctoral fellowship from the Damon-Runyon Cancer Research Foundation. M.E.V. is supported by the National Cancer Institute (NCI) of the National Institutes of Health under the Ruth L. Kirschstein National Research Service Award (F31CA257625). B.E.B. is the Richard and Nancy Lubin Family Endowed Chair at the Dana-Farber Cancer Institute and an American Cancer Society Research Professor.

## Author information

Z.C., N.J., F.J.N., and B.E.B. conceived the study. Z.C., F.J.N., and B.E.B. designed the experiments. Z.C. and M.M. performed the experiments. M.E.V. and L.P. provided computational assistance. Z.C., N.J., J.W. analyzed the data. Z.C., N.J., F.J.N., and B.E.B. interpreted the data and wrote the manuscript.

## Declare of Interests

B.E.B. declares outside interests in Fulcrum Therapeutics, HiFiBio, Arsenal Biosciences, Design Pharmaceuticals, Cell Signaling Technologies, and Chroma Medicine.

