## Supplemental Figure 1 for "Integrative dissection of gene regulatory elements at base resolution"

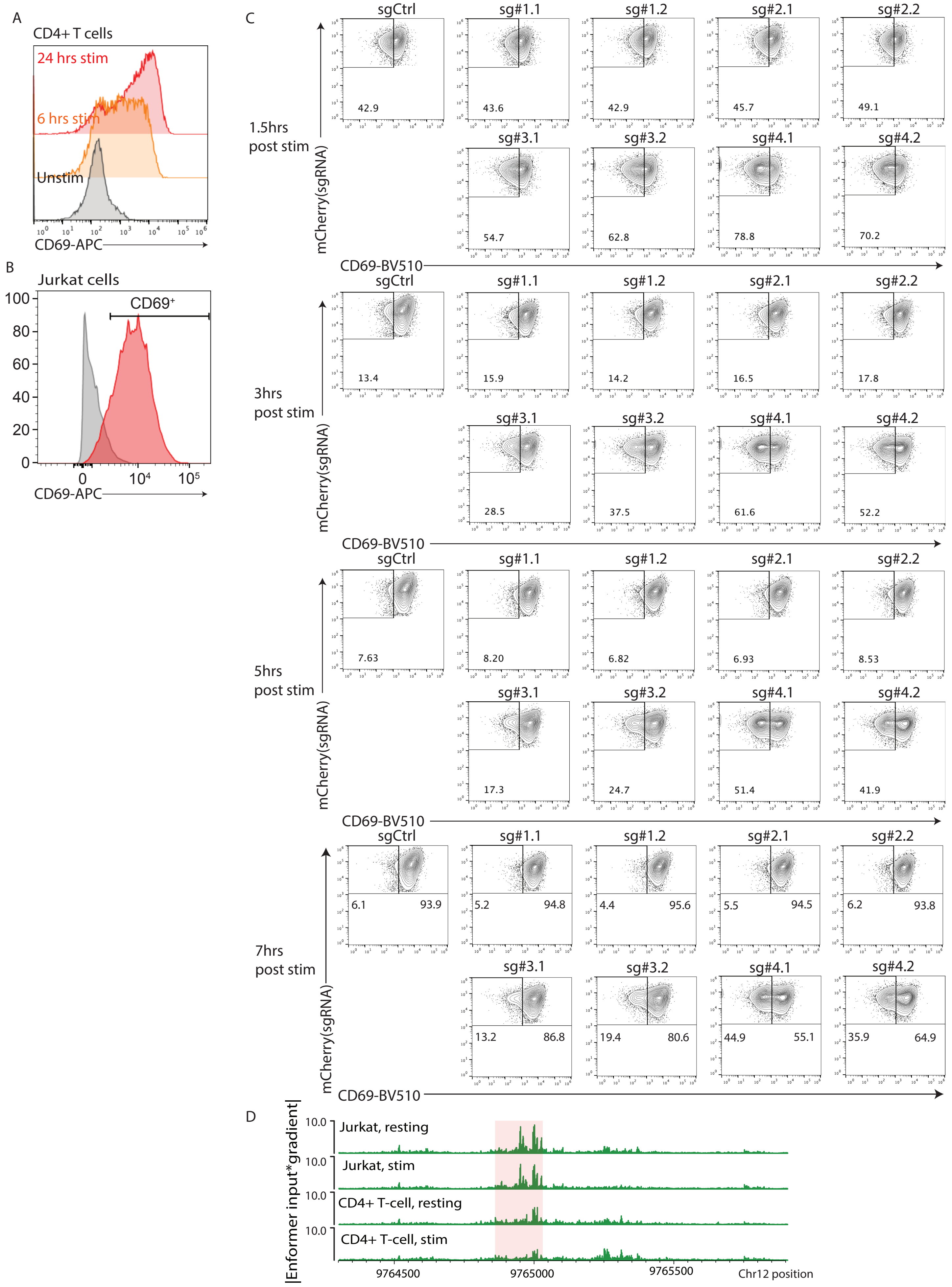


Supplementary Figure 1. Dissecting CD69 regulatory elements upon stimulation.
A) CD69 expression in primary CD4+ T cells following antiCD3/28 stimulation. Unstimulated, 6 hours stimulation and 24 hours stimulation are shown in grey, orange and red color. Data represent 2 independent experiments.
B) CD69 expression of Jurkat cells with and without stimulation. Cells were stimulated with PMA/Ionomycin for 5 hours. Data represent 3 independent experiments.
C) Flow cytometry plots of CD69 expression in Jurkat cells targeted with the indicated CRISPRi sgRNA following PMA/Ionomycin stimulation. Samples gated on mCherry+ populations. Data represent 2 independent experiments with 2 sgRNAs per targeting region.
D) Magnitude of input*gradient across RE-4 of the fine-tuned Enformer model with respect to the indicate output heads.
