## Supplemental Figure 2 for "Integrative dissection of gene regulatory elements at base resolution"

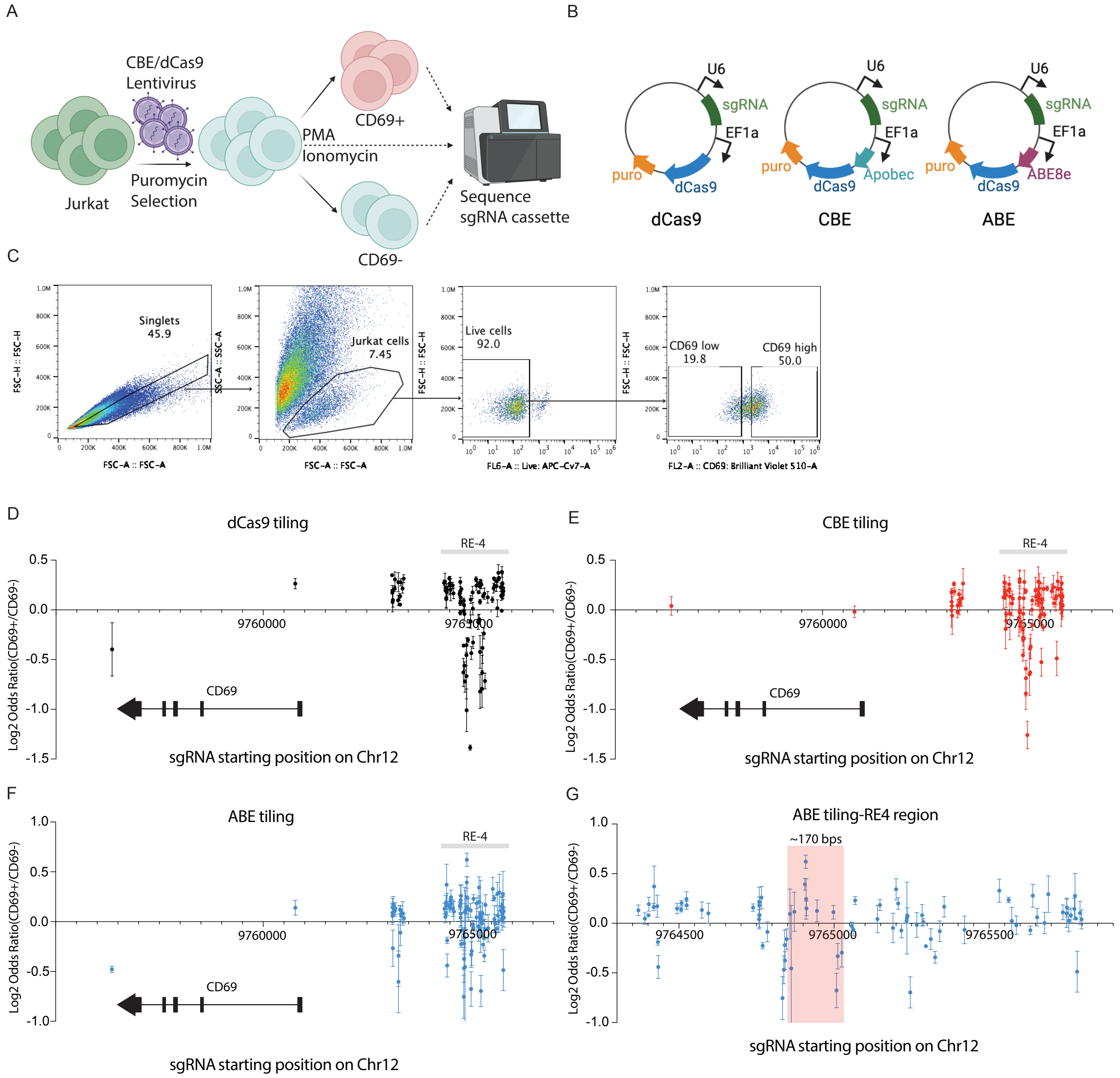


Supplementary Figure 2 dCas9, CBE and ABE tiling for CD69 loci.
A) Experimental workflow for dCas9 or CBE tiling of the CD69 loci.
B) dCas9+sgRNA, CBE+sgRNA and ABE+sgRNA vectors used for the experiment(details in Methods).
C) Example gating strategy of cytidine base editor tiling experiment. Top 50% CD69 expressed cells are sorted as CD69+; Bottom 20% CD69 expressed cells are sorted as CD69-.
D) Enrichment/depletion plot of dCas9 sgRNAs in CD69+ Jurkat cells, relative to CD69- cells (y-axis; Log2 Odds Ratio of normalized sgRNA reads). sgRNAs along the x-axis according to their 5' starting position on the positive strand. Each data point represents mean±s.e.m.
E) Enrichment/depletion plot of Cytidine Base Editor (CBE) sgRNAs in CD69+ Jurkat cells, relative to CD69- cells (as in panel D).
F-G) Enrichment/depletion plot of Adenine Base Editor (ABE) sgRNAs in CD69+ Jurkat cells, relative to CD69- cells (as in panel D) across all sgRNAs(F) or sgRNAs targeting RE-4(G). Red region shows the machine learning predicted interval.
