## Supplemental Figure 3 for "Integrative dissection of gene regulatory elements at base resolution"

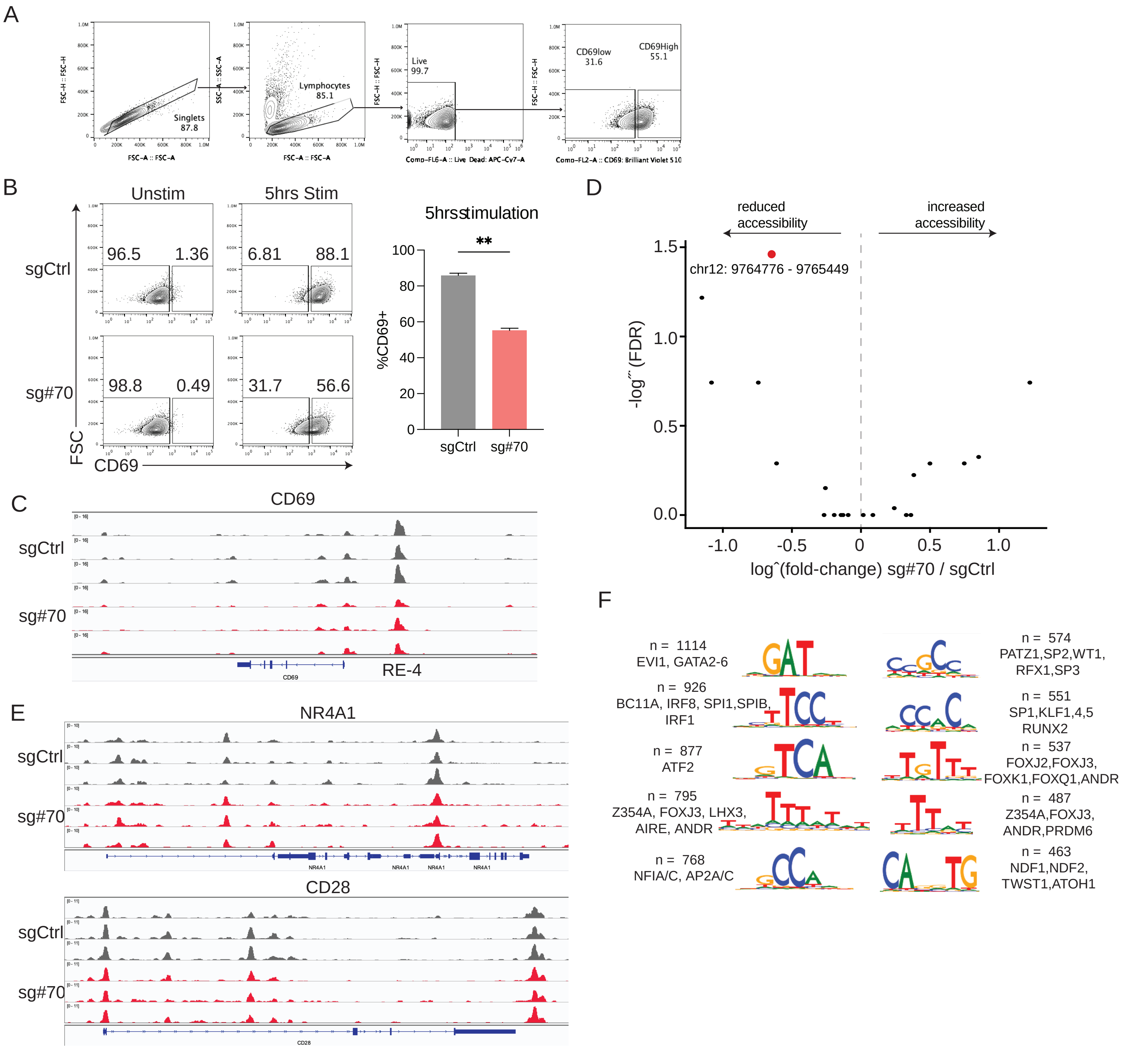


Supplementary ˜gure 3 CBE-sg#70 targeting speci˜cally a˚ects CD69 expression via RE-4.
A) Example gating strategies of CBE-sgRNA targeting experiment.
B) Flow cytometry plots of CD69 signal for CBE-sgCtrl and CBE-sg#70 Jurkat cells under resting or stimulated conditions. Bar plot depicts the propo CD69+ cells in CBE-sgCtrl (grey) and CBE-sg#70 (red) after stimulation. P-value based on unpaired t test, **P<0.01. Data are from 4 independent experiments each with 2-3 technical replicates, mean±s.e.m. Jurkat cells were stimulated for 5 hours in this panel.
C) Chromatin accessibility shown over the CD69 locus in CBE-sgCtrl (grey) and CBE-sg#70 (red) after stimulation. Biological replicates are shown.
D) Volcano plot depicts chromatin accessibility changes between sgCtrl and sg#70 groups within a 2mb window around RE4(red). X-axis shows log˛(fold-change) for sg #70 peaks relative to sgCtrl while Y-axis shows -log˝˜ (FDR), with BH correction of p-values based on changes within 1 mb window around RE-4.
E) Chromatin accessibility shown over the NR4A1 and CD28 locus in CBE-sgCtrl (grey) and CBE-sg#70 (red) after stimulation. Biological replicates
F) TFModisco identi˜ed motifs and closest TomTom matches from Enformer base importance scores. Gradients with respect to the output at the TSS for each di˚erentially expressed gene between stimulated vs. resting Jurkats were computed. Regions overlapping ATAC-peaks within resting Jurkats were provided as input to TFModisco, and the resulting seqlet clusters were matched to known motifs in HOCOMOCO. n here refers to the total number of matched seqlets, while listed motifs are signi˜cant matches from HOCOMOCO using TomTom with q < 0.05.
