## Supplemental Figure 4 for "Integrative dissection of gene regulatory elements at base resolution"

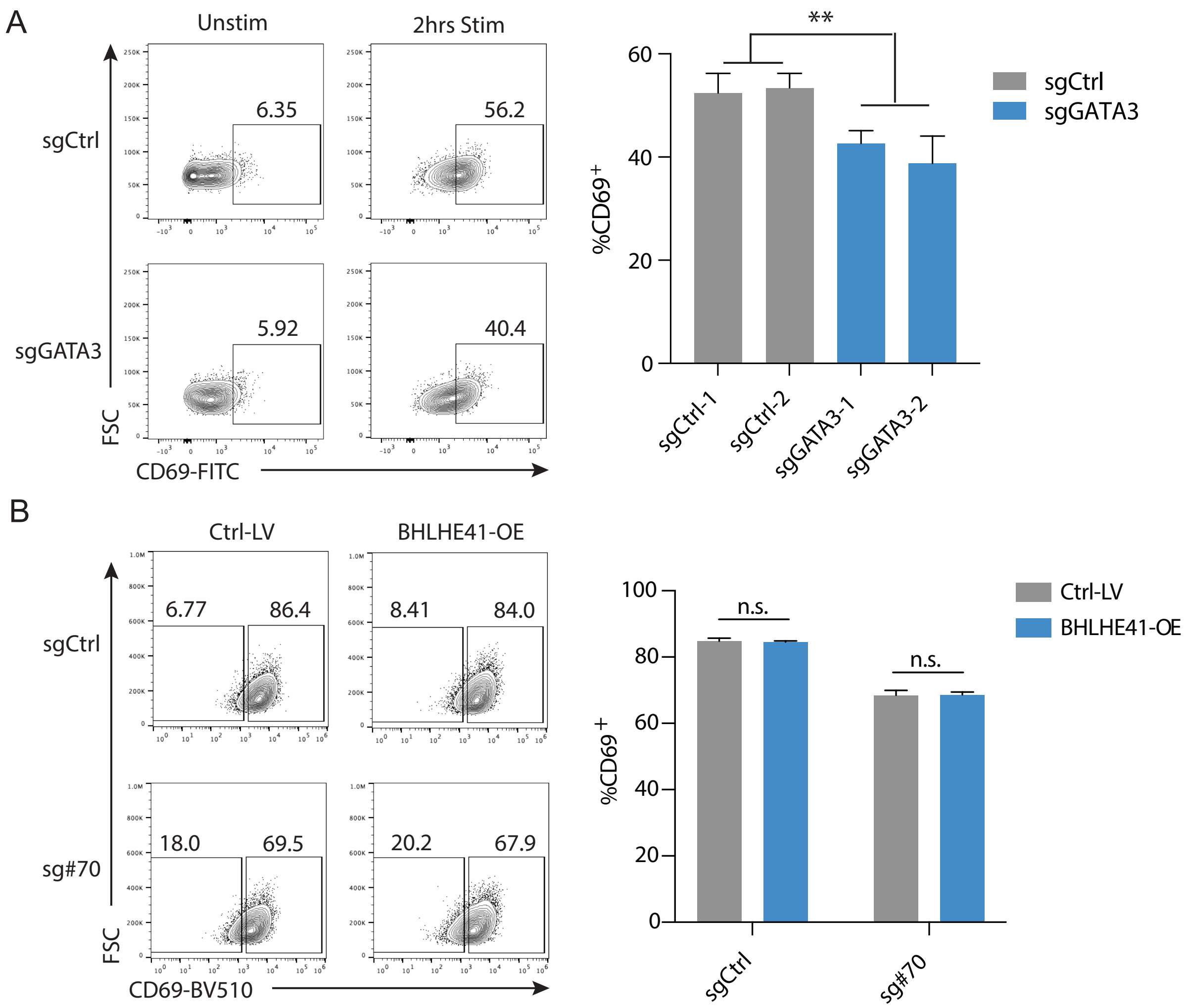


Supplementary Figure 4 Transcriptional control of CD69 expression.
A) Flow cytometry plots show CD69 expression for Cas9-sgCtrl+ or Cas9-sgGATA3+ stimulated Jurkat cells. Bar plot shows the proportion of CD69+ cells in each condition. Data represent 2 independent experiments each with 2-3 technical replicates. P-value based on unpaired t test, **P<0.01. Data are from 3 independent experiments with 2-3 technical replicates, mean±s.e.m.
B) Flow cytometry plots of CD69 signal for stimulated Jurkat cells transduced with CBE-sg#70 and a BHLHE41 overexpression construct (BHLHE41-OE), or with corresponding controls (sgCtrl and Ctrl-LV, respectively). Bar plot depicts the proportion of CD69+ cells in each condition. P-value based on unpaired t test. Data are from 2 independent experiments with 2-3 technical replicates, mean±s.e.m.
