## Supplemental Figure 5 for "Integrative dissection of gene regulatory elements at base resolution"

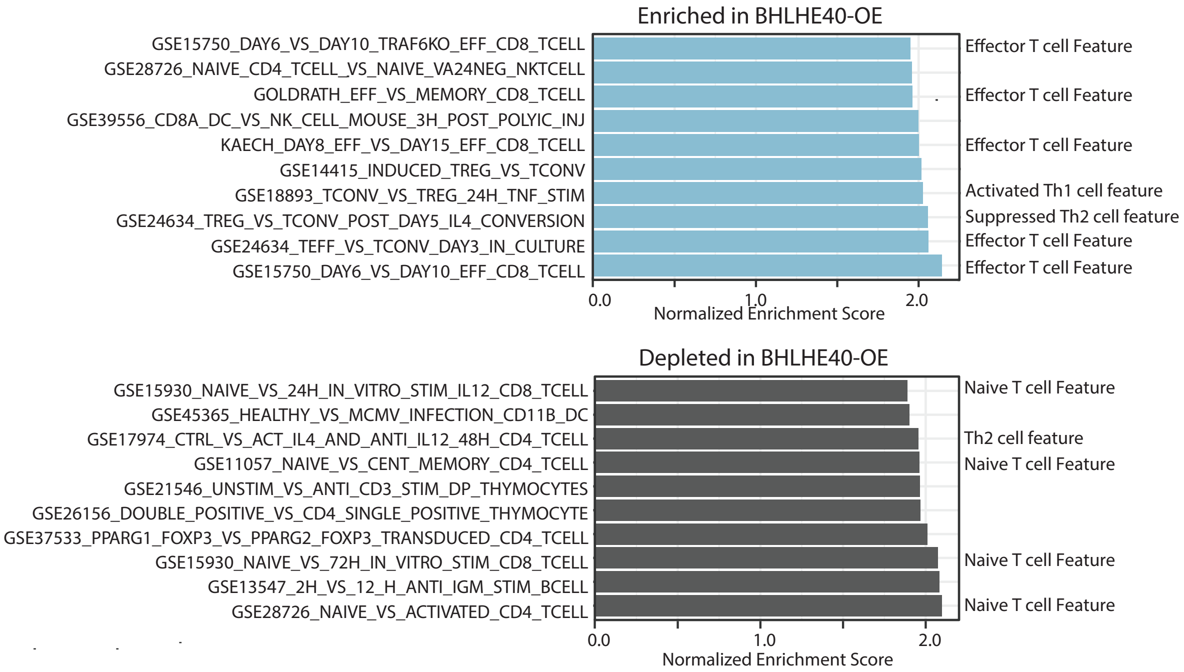


Supplementary Figure 5 BHLHE40 promotes effector T cell or Th1 cell associated features, meanwhile inhibits naive T cell or Th2 cell associated genes. Gene Set Enrichment Analysis(GSEA) in BHLHE40-WT (Ctrl-LV) and BHLHE40-OE (Overexpression) groups using RNA-Seq data. Upper panel indicates the genesets enriched in BHLHE40-OE group and the lower panel indicates the genesets depleted. Part of the GSEA feature summary was listed on the right side. X-axis shows the normalized enrichment score.
